# The multi-scale complexity of human genetic variation beyond continental groups

**DOI:** 10.1101/2024.12.11.627824

**Authors:** María J. Palma-Martínez, Yuridia S. Posadas-García, Brenda E. López-Ángeles, Claudia Quiroz-López, Anna C. F. Lewis, Kevin A. Bird, Tina Lasisi, Arslan A. Zaidi, Mashaal Sohail

## Abstract

Traditional clustering and visualization approaches in human genetics often operate under frameworks that assume inherent, discrete groupings^1,2^. These methods can inadvertently simplify multifaceted relationships, functioning to entrench the idea of typological groups^3^. We introduce a network-based pipeline and visualization tool grounded in relational thinking^4^, which constructs networks from a variety of genetic similarity metrics. We identify communities at multiple resolutions, departing from typological models of analysis and interpretation that categorize individuals into a (predefined) number of sets. We applied our pipeline to a dataset merged from the 1000 Genomes and Human Genome Diversity Project^5^, revealing the limitations of traditional groupings and capturing the complexities introduced by demographic events and evolutionary processes. This method embraces the context-specificity of genetic similarities that are salient depending on the question, markers of interest, and study individuals. Different numbers of communities are revealed depending on the resolution chosen and metric used, underscoring a fluid spectrum of genetic relationships and challenging the notion of universal categorization. We provide a web application (https://sohail-lab.shinyapps.io/GG-NC/) for interactive visualization and engagement with these intricate genetic landscapes.

## Introduction

The idea that population categories correlate with older racial categorizations traces back to the evolutionary synthesis, where the genetic concept of race was reformulated within the framework of populations. Dunn and Dobzhansky (1946) asserted that “races can be defined as populations which differ in the frequencies of some gene or genes”^6^. Although this reformulation intended to catalyze a shift from typological thinking to population thinking, it ultimately preserved the underlying assumption that human populations represent discrete, stable, natural categories. Typological concepts persisted in descriptions of human diversity, including in the UNESCO statements that retained humanity’s division into major racial categories^7,8^.

High-profile studies, such as those by Rosenberg et al. (2002)^1^ and the 1000 Genomes Project (2015)^2,5,9–11^, have played a pivotal role in structuring our comprehension of human genetic variation, and also classified populations into discrete blocks along “continental” lines (calling these super populations) for analysis and visualization purposes, further illustrating the incomplete transformation from typological constructs. Nonetheless, a growing body of literature has begun to question the efficacy and implications of such categorical classifications. Critiques by Lewis, *et al.* (2023)^12^ and the National Association of Science, Engineering, and Medicine^13^, among others, have pointed out the limitations of using continental labels as population descriptors, arguing that these categories oversimplify the rich tapestry of human genetic diversity and history. They also lead to the erroneous belief that these classifications validate a genetic basis for race^14–16^.

### Advancing beyond traditional heuristics

While this consensus against simplistic categorical labels is growing, the methods used to study genetic variation have lagged. Commonly used model-based approaches to infer population structure, such as ADMIXTURE^17^ and STRUCTURE^18^ require researchers to pre-specify the number of source populations that are assumed to be in Hardy-Weinberg Equilibrium. Model-free methods for analyzing population structure, like Principal Component Analysis (PCA), do not require a pre-specified number of categories, but are often combined with subjective approaches for identifying groups such as a sample’s continent of origin.

In response to the limitations of static, geographically defined labels, the community has sought out novel approaches and interactive tools that provide a refined understanding of genetic diversity. From the Geography of Genetic Variants (GGV) browser to the “Visualizing human genetic diversity” blog, and employing methodologies like FineStructure, topological analysis, and ancestral recombination graphs, researchers are exploring genetic variation in richer ways that challenge traditional views^19–24^. Despite these shifts, STRUCTURE/PCA are still dominant.

### Our Contribution: Contextual and Fluid Groupings

In biomedicine, genetic similarity is now widely understood to be more relevant than (continental) ancestry for interpreting and accounting for genetic structure^13^. Network approaches have recently emerged as fruitful for decoding the genetic structures that may underpin disease risk and other aspects of human health^22,25–29^. Network-based approaches capture complex relationships among individuals with minimal assumptions and without need for a pre-specified number of populations. Further, a suite of established community detection algorithms can identify subnetworks called communities, grouping genetically similar individuals. Communities are always connected internally, as well as externally to individuals in other communities. They are also fluid in the sense that their composition and subsequent connections vary depending on the resolution considered. Previous implementations of network analyses have primarily focused on specific aspects, like demonstrating the feasibility of network approaches^26,28^, or leveraging networks to identify hierarchical structure^25^.

We present a novel framework called the Global Genetic Network Communities pipeline and browser. It centers two key aspects of network analysis. First, it allows for great flexibility in the definition of genetic similarity, both the metric used and which data are used to compute it. Second, the detection of communities at varying resolutions can be achieved using any suitable community detection method. Both of these aspects facilitate more dynamic data-driven, assumption-free analyses and visualizations of genetic structure suited to the particular questions targeted in a study. These groupings convey the landscape of genetic diversity without fixed or geographically bound labels, thereby challenging oversimplified classifications and fostering a more interconnected and fluid view of genetic diversity that aligns with the realities of human evolution and migration.

## Results

### Flexible community detection with GG-NC

Our Global Genetic Network Communities (GG-NC) pipeline accepts diverse genetic similarity metrics (Figure 1 and S1) to construct networks which represent individuals’ genomes as nodes and genetic similarity as edge weights. Genetic similarity can be defined in different ways (e.g. identity-by-descent sharing, and kinship), with different sets of variants (e.g. common vs rare), or based on different parts of the genome (genome, exome, or trait-specific variants), allowing users flexibility in probing genetic similarity as a function of evolutionary timescale and functional importance. Our pipeline uses the Louvain algorithm^30,31^ to infer modules or communities in these networks at different resolutions^25,32^. However, we also implement the Leiden algorithm, which has some superior properties^33^ (see methods).

**Figure 1.**
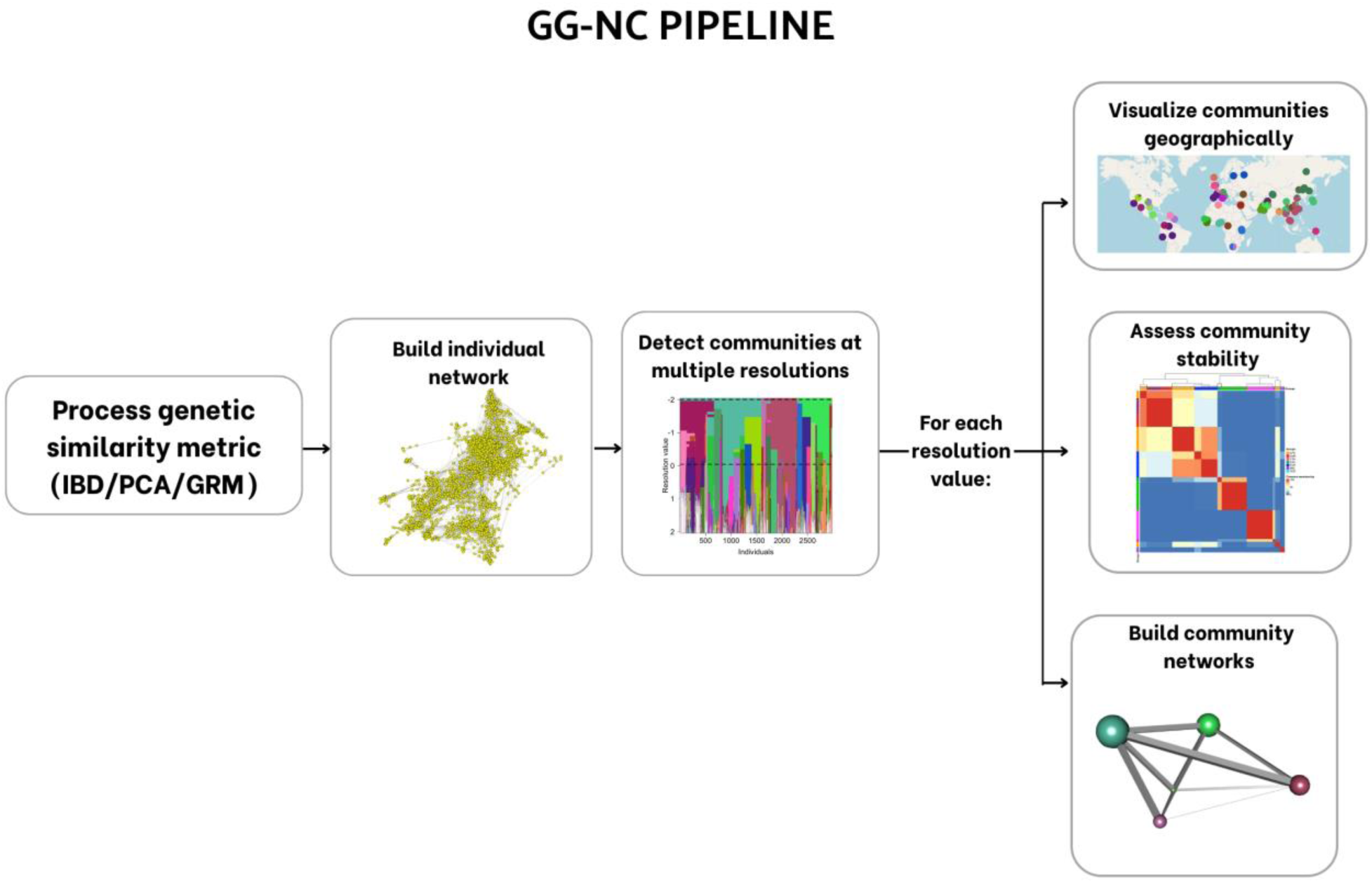
Overview of the Global Genetic Network Communities (GG-NC) computational pipeline. GG-NC is grounded in relational thinking in contrast to typological thinking^4^. Communities are detected on genetic similarity networks at multiple resolutions. Across these resolution values, our pipeline computes the stability of the detected communities, builds networks of the detected communities, and visualizes the detected communities geographically on a world map.

No single metric or resolution value is considered correct; instead, we explore the effect of the parameter space on the communities detected using *resolution plots* (Figure 2), which summarize the communities detected across a range of resolutions. In a *resolution plot*, each vertical line represents the same individual allowing us to observe their changing community membership and how communities break apart into smaller ones as the resolution value is increased.

**Figure 2.**
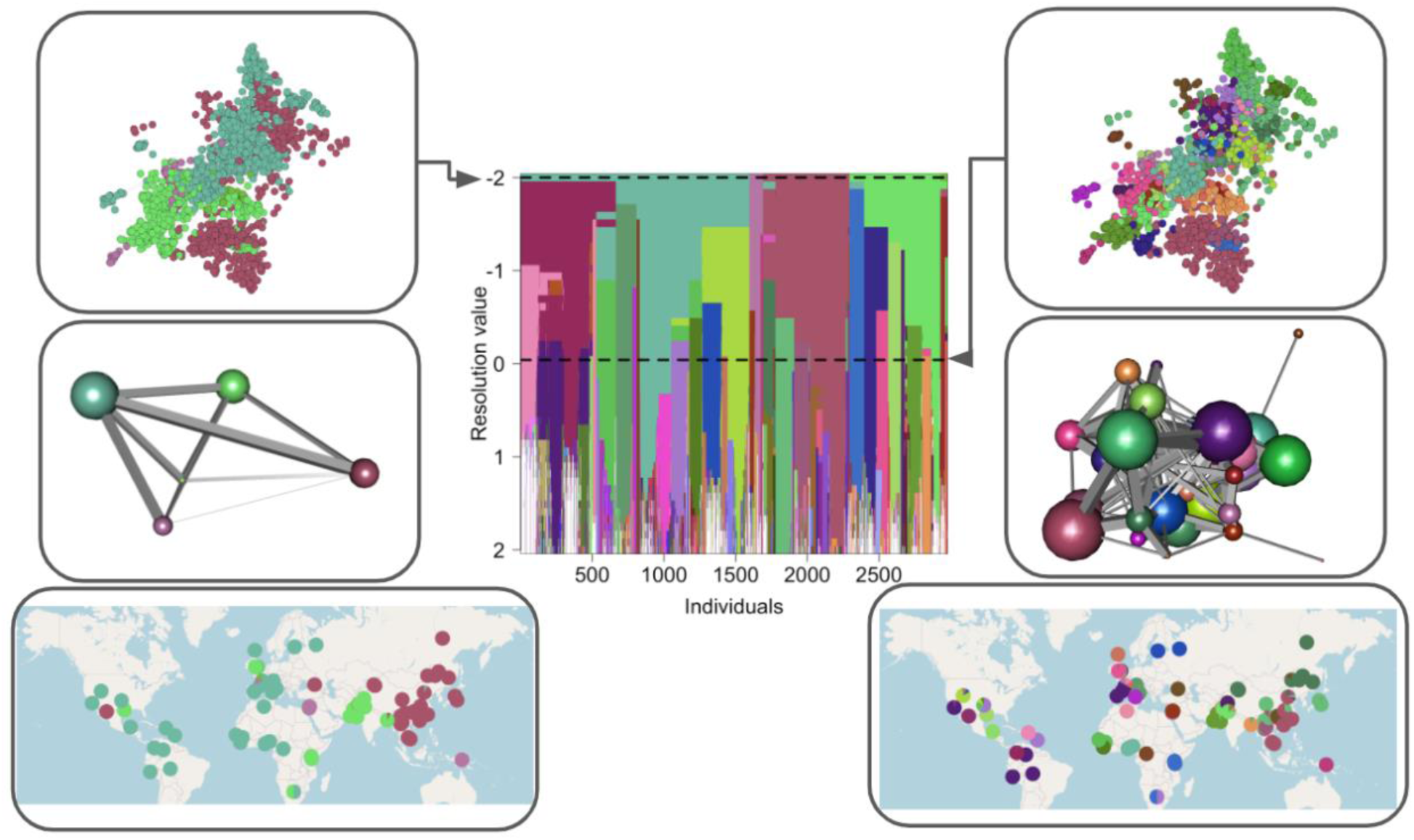
*Resolution plot* for IBD results and associated individual and community networks, along with the geographic distribution of the communities. The central plot is the *Resolution plot* showing the results of the Louvain algorithm at 50 resolution values (see results for Leiden algorithm in Figure S3). Communities that do not have more than 6 members in any resolution are colored in white. The left panels display results for a resolution value of −2, and the right panel shows results at resolution −0.041. Each side includes (from top to bottom) the individual network, the community network, and the geographic distribution of the communities. The individual network is formed of 2,977 individuals represented by nodes (280 outlier samples were excluded from the network for visualization purposes, see methods and Figure S4) in which nodes are colored according to the community membership in the *resolution plot*. Community network plots present communities as nodes and the density of the connection among them as edges. In the maps, we show the 1000G project and HGDP cohorts using pie charts placed at sampling locations. Each pie chart represents the community membership of the individuals within each cohort. Finally, a color-coding scheme was implemented where genetically “closer” communities are represented by more similar colors (see methods and Figures S5 and S6).

There is an inherent stochasticity to community detection algorithms, and therefore, the community that each individual belongs to at a given resolution may shift across different runs. To allow users to assess the stability of the communities detected at a given resolution value, the pipeline computes the Adjusted Rand Index (ARI) and Normalized Information Distance (NID) (see methods).

Once communities are detected, we emphasize the continuum of relationships among them by creating community networks that represent communities as single nodes and the density of the connection between them as edges, with the size of the nodes being proportional to the size of the community.

Finally, we visualize the geographic distribution of the detected communities at multiple resolutions using a web browser that we developed. Our results are available through the browser (https://sohail-lab.shinyapps.io/GG-NC/) and our computational pipeline is flexible and accessible, allowing extensions to any new dataset (https://github.com/mariajpalma/GG-NC).

To illustrate our approach, we computed the pairwise genetic similarity among 4,150 individuals from the harmonized 1000 genomes project and Human Genome Diversity Project (HGDP) dataset^5^ using four different metrics: (i) the Genetic Relationship Matrix (GRM) using rare variants (ii) GRM using common variants, (iii) Correlation of PC scores (PC), and (iv) sharing of identity-by-descent (IBD) segments (Figure S2 and Supplementary Tables 1-3). The use of these different inputs enables us to probe genetic similarity at different evolutionary timescales^34,35^.

### Genetic communities beyond continental groups

Our approach allows the user to examine groups from multiple “viewpoints” providing insights into genetic structure that is highly dynamic. Our results show that there is no clear basis to structure individuals in genetic studies primarily by continental origin. We demonstrate this in Figures 2, 3, and 4 using community detection on sharing of IBD segments longer than 5cM which is useful in studying recent demographic history and fine-scale genetic structure^34^ (results from other metrics in Figure 5 and the supplement (Figures S7 and S8). At a low resolution value of −2, representing a “zoomed out” view of the network structure, five major communities emerge (Figure 2). The largest comprises 1595 individuals (shown in teal) with a wide geographic distribution including individuals from the Americas, Europe, and Africa. The other two communities are colored in deep rose and bright green respectively with around 600 members each. The deep rose community includes individuals from East Asia and Pima individuals in Mexico while the bright green community is mainly formed by individuals from Central South Asia, including Gujarati Indians in Texas, Indian Telugu in the UK, and Sri Lankans in the UK. A community with 95 individuals (colored in orchid) is formed of Palestinians, Bedouins, Papuans, and some French individuals. The smallest community of 50 individuals (shown in green grass) groups together Hazara and Druze individuals. Community networks show the relationships that exist among these communities, for instance, showing a closer relationship between bright green and teal communities than the bright green and deep rose communities.

**Figure 3.**
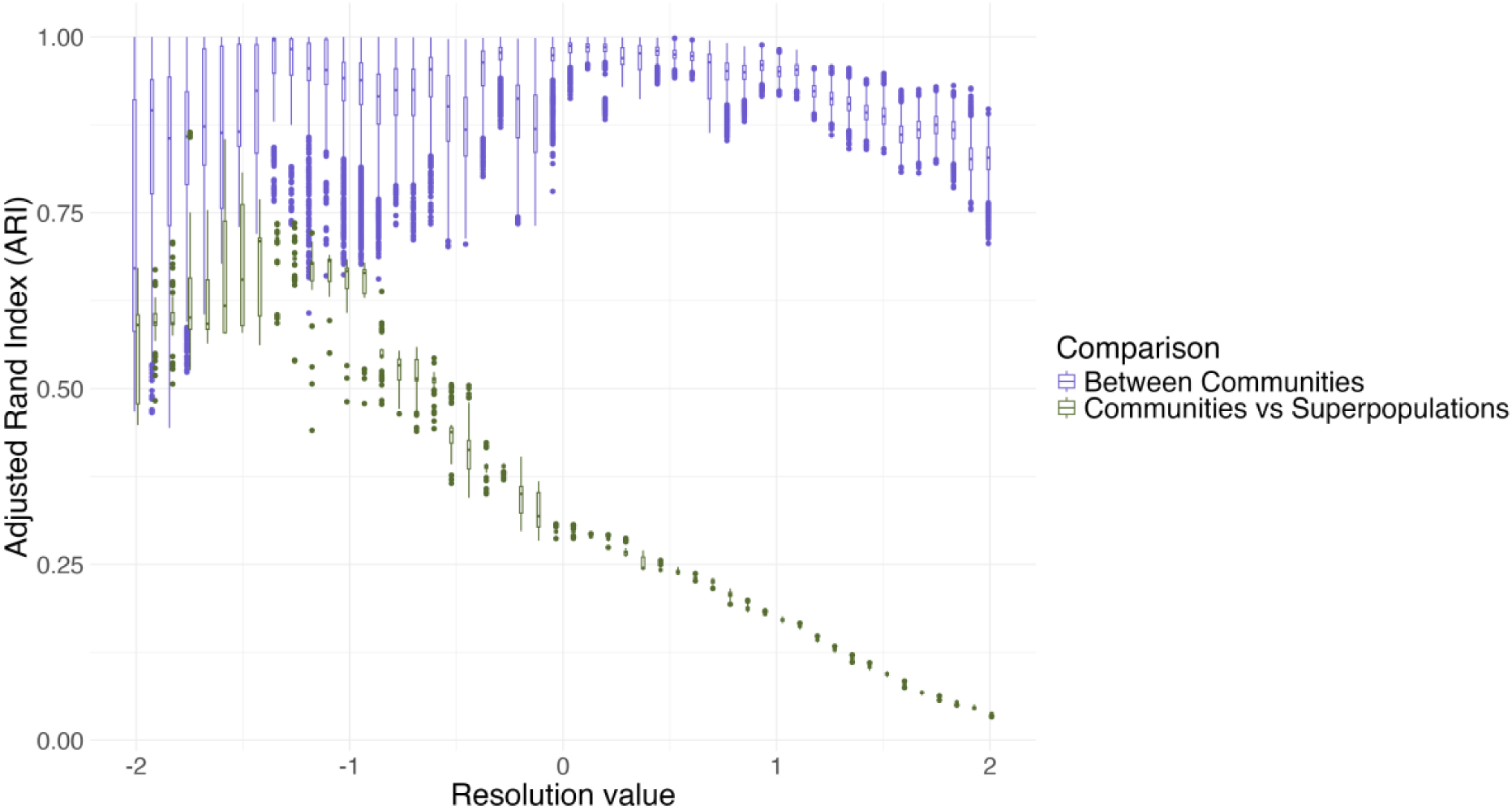
Communities detected in the IBD-network are fairly stable across resolutions, and different from superpopulations from 1000G and HGDP. The x-axis shows the resolution value. The y-axis shows the ARI values. ARI values closer to 1 indicate more individuals falling in the same communities across runs at a given resolution value. Purple boxplots summarize the comparison of community detection results across 100 independent runs at each resolution (see methods). Green boxplots represent the comparison between the independent runs and the super populations. In this case, ARI values closer to one indicate greater similarity between the detected communities and the superpopulations. Boxplot elements: center line, median; box limits, upper and lower quartiles; whiskers, 1.58x interquartile range; points, outliers. The same analysis was conducted for GRM and PCA networks (supplementary figures S12 and S13 (NID)).

**Figure 4.**
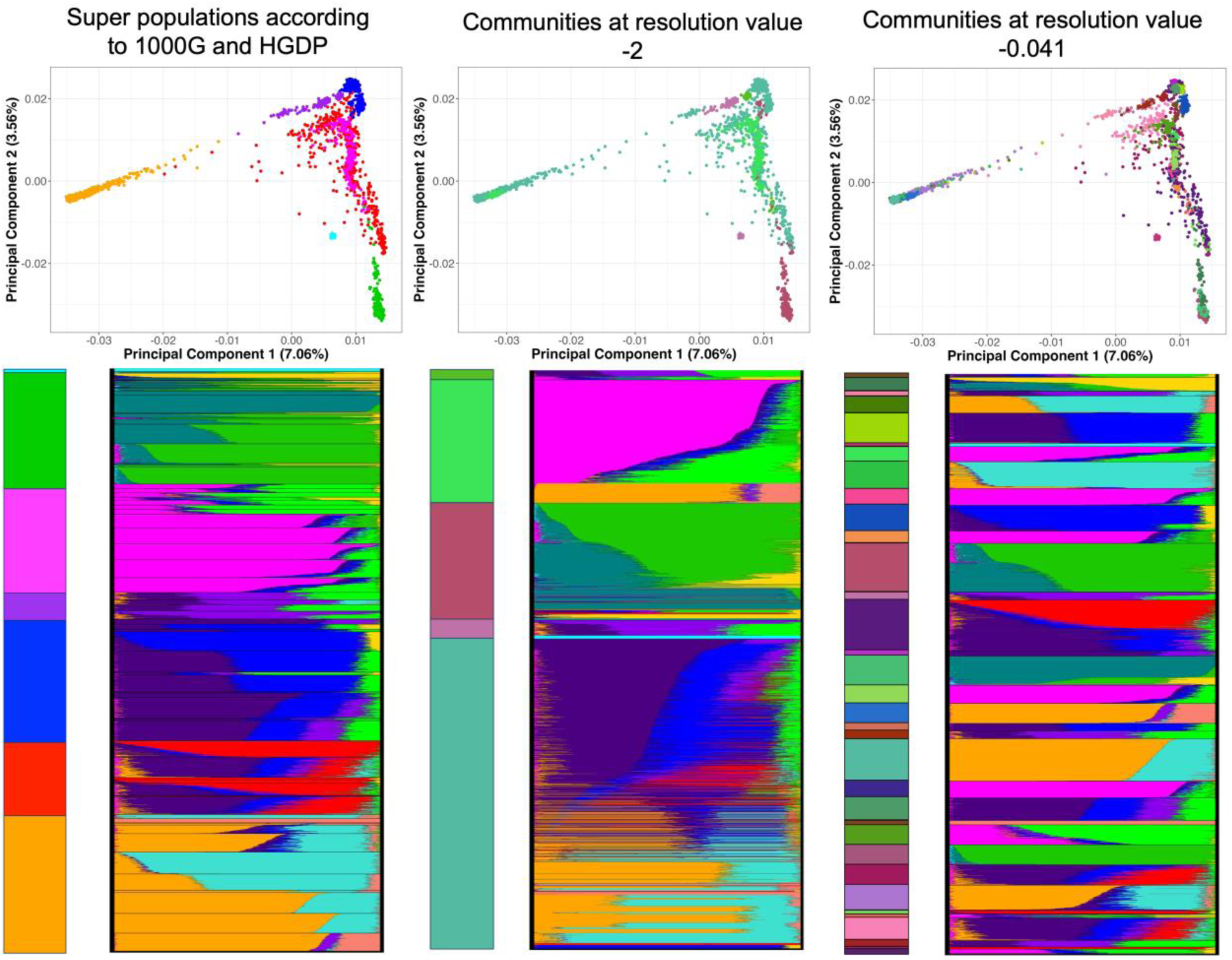
Using communities derived from GG-NC gives different insights than conventional population and super population groupings. Each column shows the same type of information but using different groups illustrated with different colors. In column A, the colors come from the standard super populations (7 groups; Supplementary Table 2). In columns B and C, they come from the communities detected at different resolution levels: −2, where 5 communities are detected, and at −0.041 where 34 communities are detected. At the top of each column is a PCA plot, created from the jointly called dataset of 1000G and HGDP (2,977 samples included in the shown networks). At the bottom right of each column is an ADMIXTURE plot using the same data and K = 13 (lowest cross-validation error), but with individuals sorted by the different color grouping, according to the stacked bar chart at bottom left. Community membership at the two different resolutions gives different insights than the conventionally deployed superpopulations.

**Figure 5.**
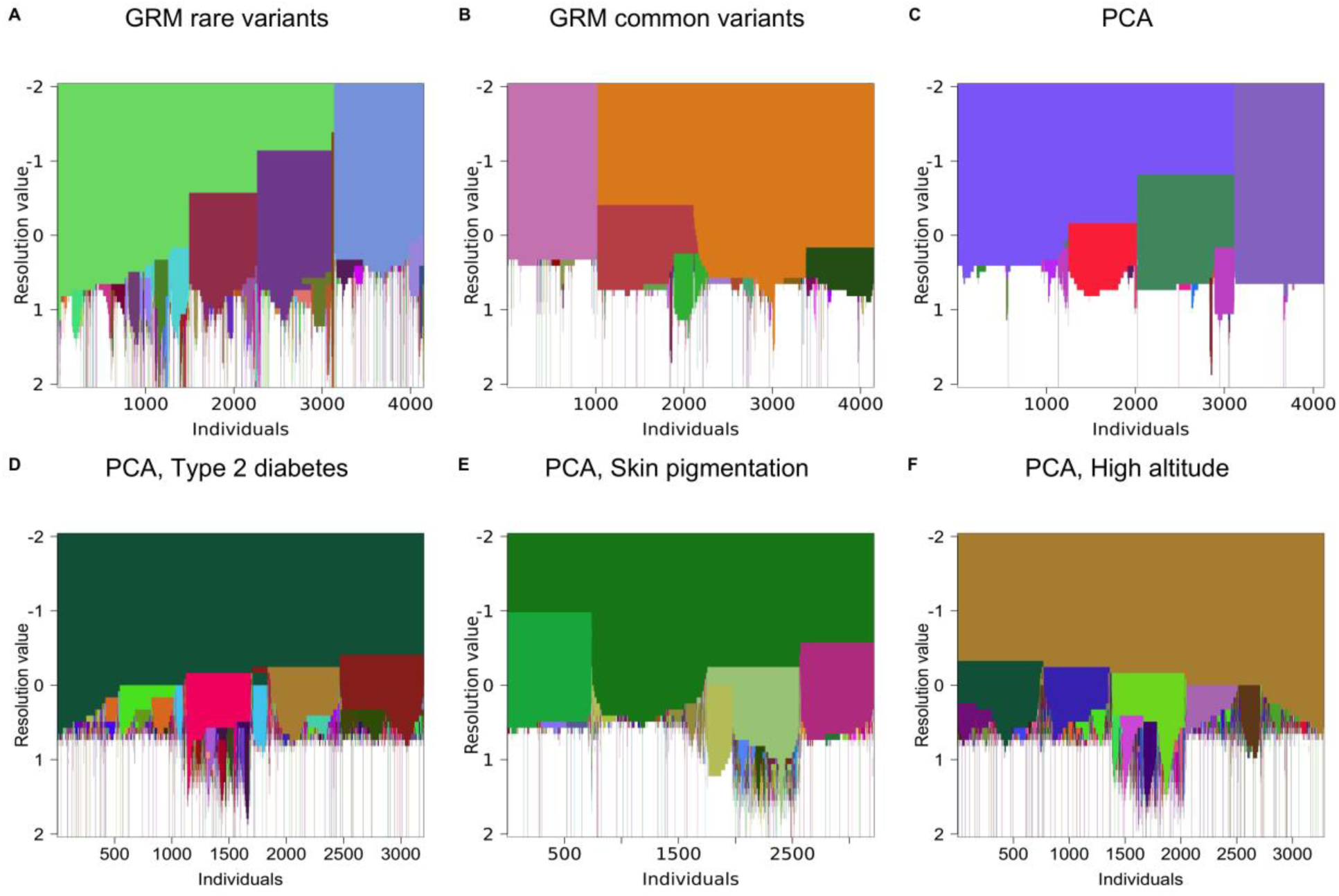
*Resolution plots* from networks using different definitions of genetic similarity and different subsets of genetic variants reveal different aspects of genetic relatedness. *Resolution plots* summarize community detection results at 50 resolution values; Communities that do not have more than 6 members in any resolution are colored in white. The x-axis represents the individuals and the y-axis corresponds to the resolution value (see results for Leiden algorithm in Figure S20). A) *Resolution plot* for the network based on the Genetic Relationship Matrix (GRM) estimated on rare variants (n=4,150). B) *Resolution plot* for the network based on the GRM estimated on common variants (n=4,150). C) *Resolution plot* for the network based on Principal Component Analysis (PCA) correlation (n=4,119). *Resolution plots* for trait-PCA-based networks using only independent variants in: D) Type 2 diabetes associated genes (n= 3,199; 15 genes)^36^. E) Skin pigmentation associated genes (n=3,214; 38 genes)^37^. F) Genes associated with or inferred to be under natural selection for Altitude adaptation (n=3,281; 7 genes)^38^.

At a higher resolution of −0.041, the number of communities increases to 34 (Figure 2). Some cohorts from geographically close regions such as Pima and Maya indigenous groups in Mexico form distinct communities with each other. In contrast, individuals from different continental groups remain in the same community such as Mexicans in Los Angeles, Peruvians in Lima, Colombians in Colombia, Karitinian in Brazil, Iberian populations in Spain, Basque in France, and French in France. Importantly, clear substructures appear within continental groups, even before continental groups split from each other. For example, individuals from Africa are grouped into 7 communities. Afro-descendant individuals in the Americas are grouped with four of these communities showing the diversity of the genetic ancestries that contributed to the Afro-descendant groups. Substructure within countries is also evident, for example, individuals in Pakistan are mainly grouped into 4 communities: Makrani, Barahui, and the majority of Sindhi and Balochi individuals (as well as a few Pathan and Punjabi individuals) belong to one of these communities, all Hazara individuals are part of a different community along with a few French individuals, Burusho individuals form the third community, and the majority of Punjabi and Pathan individuals are grouped into the fourth community.

We further ask how stable our communities are at each detected resolution, and how they compare to the standard continental labels used in many human population genetic studies. To answer this, we estimated the pairwise Adjusted Rand Index (ARI) value for 100 replicates to assess “stability” of detected communities at every resolution (Figure 3, see Figures S9 and S10, supplementary text and methods). We show that at lower resolutions, the median ARI of the communities detected through the GG-NC pipeline is low with a high variance, suggesting that community membership is highly unstable across runs even though some individuals might be consistently grouped together in the same community. Stability increases at higher resolution values, peaking after R=0 with a low variance, suggesting more consistent grouping of individuals, before decreasing again slightly at higher resolution values where more and more communities are observed. We also formally compare continental “super population” labels and the communities detected in the network across all resolution values for individuals from the HGDP and 1000G datasets, answering some key questions about their correspondence. Is there a point where continental labels are equivalent to the network communities? No, at every resolution, the communities identified on the IBD network differ from the superpopulations of the 1000 Genomes Project and HGDP (median ARI communities vs. super population <= 0.71). Even at the resolution (R=-1.34) where we observe the highest concordance between super populations and communities detected (median ARI = 0.71), the variance of both ARI distributions is large suggesting a lack of consistency in community membership, and we detected 12-14 communities using GG-NC compared to only 7 superpopulations (Figure S11). Are the network communities detected similar to continental groups at a majority of resolution values? No, at resolutions greater than −1, the similarity between super populations and network communities decreases linearly. In fact, the network communities are more stable amongst themselves than they are with super populations at every resolution (Supplementary Table 4, Wilcoxon test). Given this, the standard use of continental groups to organize or visualize individuals in genetic studies seems poorly suited if the goal is to accurately and faithfully represent patterns of genetic similarity. Instead, the communities detected based on genetic relationships transcend continental boundaries at low and high resolutions.

### Comparisons to traditional approaches

A comparison of our approach with existing approaches such as ADMIXTURE and PCA further illustrates the dynamic complexity of human genetic variation. In particular, IBD-based data-driven clustering does not recapitulate the clean super-populations that the 1000 Genomes and HGDP studies have used to frame human genetic variation. To show this, we carried out ADMIXTURE (K = 13 with the lowest cross-validation error) and PCA (first 20 PCs) on the same dataset (n=2,977; Figure 4, Figures S14-16). First, we grouped individuals according to pre-defined continental categories (super populations) from the 1000G and HGDP studies, and colored PCA results according to these continental labels (Figure 4A). Alternatively, individuals in the admixture plot were grouped according to the 5 communities detected at resolution value −2 (Figure 4B), and the 34 communities found at resolution value −0.041 (Figure 4C) using the GG-NC pipeline based on IBD data.

The network community-based analysis reveals many levels of structure. For example, we observe that individuals from the Middle East (purple color in Figure 4A) are split into three different communities (Figure 4B) at a resolution of −2. These communities also include individuals from other continental groups (Figure S17). A distinct substructure is seen when increasing the resolution to −0.041, with individuals belonging to the same community nevertheless clustering closely in PC space (Figures 4C and S18).

Furthermore, we show that there is no direct relationship between genetic similarity (reflected by the IBD-based communities) and ADMIXTURE components. We observe individuals with different ADMIXTURE components grouped within the same community, as seen in the dark purple community (Figures 4C and S19). This community includes individuals from diverse cohorts, such as the Iberian Population in Spain, individuals with Mexican ancestry in Los Angeles, Basque in France, and Peruvians in Lima, among others.

Conversely, distinctive communities exhibit similar proportions of ADMIXTURE components. For instance, Lime green, Bright pink, and Orange communities at resolution value −0.041 share similar proportions of these components (Figures 4C and S19). These communities also occupy similar or overlapping positions in PC space (Figure 4C). The lime green community is predominantly composed of Gujarati Indiviand in Houston, while the pink community is primarily formed from Indian Telugu in the UK, and the orange community is composed of Bengali individuals in Bangladesh. Another example is that individuals with a high proportion of the red Admixture component are distributed in three different communities (Figures 4C and S19).

We show how network-based community detection captures genetic similarities that transcend the sharing of ancestry proxies as captured through an ADMIXTURE approach, primarily highlighting that communities emerging from population-based thinking (GG-NC) do not neatly fall into continental ancestry categories. A key issue/limitation with standard ADMIXTURE approaches is that they assume the existence of otherwise “pure” types. Thus, they remain confined to a typological or continental framework.

### Metric choice impacts detected structure

Another layer of complexity in the inference and visualization of human genetic diversity is the information used to specify genetic similarity. To illustrate, we applied our community detection pipeline using different measures of genetic similarity (Figure 5). We show that the number and size of communities detected are determined both by the input genetic metric and the subset of variants used. For example, in the *resolution plot* based on GRM using common variants (GRM common) (Figure 5B), we observe fewer larger communities before they eventually fragment into many smaller ones (<6 members), a pattern also observed in the *resolution plot* based on PCA. In general PCA and GRM common produce more communities at higher resolutions; however the size of these communities (<6) limits their utility for analysis and reflects an abrupt fragmentation of larger groups. In contrast, in the *resolution plot* based on GRM using rare variants (GRM rare) (Figure 5A), we observe a greater number of intermediate-sized communities, which better capture finer genetic structure (Figure S21). This generally makes sense since rare variants are more recent in origin and therefore, more useful for the study of fine-scale structure than common variants which are older in origin^35^.

Furthermore the *resolution plots* for PCA, GRM common, and GRM rare all show that at the lowest explored resolution value, two distinct communities emerge, separating Sub Saharan African individuals from the rest of the human groups. The first three communities detected on the network generated from PCA and GRM common are almost identical, dividing individuals into three major geographic areas: Sub-Saharan Africa (including Afrodecendent individuals), Europe and Central South Asia (excluding some Hazara individuals), and East Asia and the Americas. In contrast, in the GRM rare network, a community composed of individuals from Oceania is detected after the division of Sub-Saharan Africa and the rest of the world (Figure S21). This community is first detected at higher resolutions of 0.531 and 0.612 using GRM common and PCA networks respectively.

While increasing the resolution value, a high proportion of Hazara individuals from Pakistan and Uygur individuals from China form their own community when analyzing GRM common (R=0.449) (Figure S22) and PCA networks (R=0.531) (Figure S23). Hazara and Uygur individuals are also grouped when analyzing GRM rare networks, along with individuals sampled in China (Xibo, Mongolian, Oroquen, Daur, Hezheh) and Yakut individuals in Siberia. These findings corroborate previous studies on Hazara and Uygur being genetically close^39^. Despite the similarities among PCA and GRM common results, some communities such as Bedui in Negev and Druzel in Camel were detected by PCA and GRM rare networks, but not in the GRM common network.

Further, not all genes or regions of the genome reflect the same evolutionary history; therefore the genetic similarity of individuals will not be identical for all loci. We highlight that the relevant communities for a gene or a given set of genes (related to a phenotype of interest) may differ from one set of genes to another. The groupings most relevant for genetic epidemiology depend on the specific sets of genetic loci and the trait under consideration. To demonstrate this, we analyze sets of specific genes involved in Type 2 Diabetes, skin pigmentation, or altitude adaptation at diverse resolutions using PCA-based networks (Figure 5, d-f, Supplementary text, Supplementary Figures S24-S27 and Extended Data Tables 1-3). Similar to Mohsen et al^40^, we believe that this approach can allow users to explore the community structure relevant for genetic variation associated with their trait of interest, to help identify trait-specific variant clustering and epidemiology that may not relate to continental categories.

We tested and showed that, as for the IBD networks, network communities detected from any of these metrics or set of variants are more stable amongst themselves than they are with super populations at every resolution (Figures S12-13, S28, and S29, Supplementary Tables 5-10, Wilcoxon test). Further, their concordance with super populations varies significantly over the resolution range. As would be expected, networks based on PCA on all variants, or on PCA on variants associated with skin pigmentation give the highest concordance with super populations at their peak value compared to other networks (Figures S11 and S30; maximum median ARI(genome-wide PCA)=0.904 and maximum median ARI(PCA on skin pigmentation associated variants)=0.887 for comparison of network communities with super populations). This makes sense as, (1) PCA on common variants best captures broad-scale patterns of variation especially when combined with sparse sampling as in the 1000 Genomes and HGDP joint dataset, whereas IBD or GRM-rare networks capture more fine-scale structure, and (2) race as a social construct was primarily created based on skin color^41^. Nevertheless, even at the resolution of their maximum concordance with super populations, the network based on PCA on common variants results in 8-11 communities, and the network based on PCA on skin pigmentation variants results in 10-12 communities, in comparison to 7 super populations. Overall, this work reinforces the idea that the genetic similarity between two individuals can be measured in different ways capturing different aspects of genomic variation, and that any scheme to cluster individuals based on genetic similarity including for biomedical purposes must take this into account.

## Discussion

Our network-based approach captures and reflects the fact that there are no universally valid or relevant groupings of genetic variation. When different genetic similarity metrics are used (e.g. IBD, rare-GRM, common-GRM, and PCA), each contains unique patterns of genetic relatedness that were not well-captured by either traditional continental divisions or standard approaches like ADMIXTURE or PCA. Our analysis of network communities based on trait-related variants further underlined that no single representation of human genetic ancestry captures genetic patterns relevant for all traits. Collectively, our study supports a shift away from traditional typologies towards a fluid, context-specific understanding of genetic diversity. Instead of viewing genetic groups as static descriptors of the world, our findings argue for an approach where decisions on how to represent genetic relationships and groups are shaped by the particular context and purpose of a study^42^.

Beyond simply challenging the use of conventional genetic groupings, our contribution is the flexibility of the GG-NC pipeline enabling multiple operationalizations of genetic similarity by using networks defined i) using any number of similarity metrics, ii) on different subsets of genetic data (e.g. just constrained to relevant to specific traits) and iii) probing these networks at multiple resolutions.

GG-NC will be useful as the starting point for research projects in genetic history or biomedicine. Researchers can use the GG-NC pipeline to quantitatively and qualitatively analyze and visualize the genetic structure in their dataset at different resolutions, and obtain graphics summarizing the multi-scale complexity of genetic variation in their dataset. They can also obtain quantitative measures of the stability of the genetic structure at any given resolution using a specific similarity metric of choice. In this way, the user can navigate different evolutionary timescales to view genetic structure from multiple “viewpoints” with ease and flexibility, before deciding upon a particular metric or resolution relevant to their question. GG-NC allows researchers to analyze the genetic structure of study samples on their own or in combination with reference datasets (e.g. 1000 Genomes and HGDP, or other cohorts sampled at finer-scales), which can be useful in studying genetic ancestries when detailed demographic information is not available. GG-NC will determine the reference individuals that study samples cluster with at different resolutions, and allow communities for specific research questions to be identified. Instead of using ancestry, continental labels, or ad-hoc clusters, we affirm that researchers should describe the genetic structure of their study samples at different resolutions and provide a justification for why they have chosen to use a particular resolution value. Future work should assess applications of GG-NC to study genetic structure in other organisms, as well as undertake theoretical analyses to relate resolutions for different similarity metrics to evolutionary timescales.

GG-NC can further serve projects interested in detecting genetic variants that are highly differentiated across groups due to selection, demographic events, or/and association with a disease or trait. In this case, researchers can use the pipeline to determine the relevant clusters, which can then be used as the unit/population for selection analysis, for example, with population branch statistics^43^, or as cohorts for association analysis that can then be meta-analyzed. The inferred communities can also simply be used to understand trait/disease variation among different communities^44^ or/and assess underlying SNP differentiation. This would have clear value for public health and precision medicine, without the need to resort to continental groups. Notably, the GG-NC enables researchers to analyze genetic structure at varying resolutions with ease, allowing one to understand at which scale the genetic community structure became relevant for a particular disease or SNP differentiation, and helping researchers to identify communities that share or carry unique genetic risk for a given disease or trait^44^.

The GG-NC pipeline and browser also provide an important educational resource that can be used in courses and workshops. Further, it is a resource that the public can use to develop an understanding of genetic diversity. In these ways, it can be a tool against white supremacists and their weaponization of genetic science towards a racist agenda.

Our approach enables researchers and the general public to shift to a more accurate, non-essentialist perspective on human diversity. It provides new tools and terminologies to foster more insightful, ethical, and inclusive explorations of our shared humanity and the relevance of genetic variation to our lives.

## Materials and Methods

### Dataset

We applied our pipeline to the recently published jointly called reference panel of the 1000 Genomes (1KGP) and HGDP projects^5^. We downloaded the set of variants jointly called on the HGDP+1KGP data and the metadata information from gnomAD (https://gnomad.broadinstitute.org/downloads#v3-hgdp-1kg) server into our HPC Kayab server. Sampling locations were obtained from https://www.internationalgenome.org/data-portal/population.

### Global Genetic Network Communities Pipeline

#### Building Individual network

We developed a computational pipeline in R (Figure 1 and S1) that uses the package igraph^45^ to build a network from an adjacency matrix or directly from a data frame. The matrices can be obtained directly from the Genetic Relationship Matrix (GRM), from the pairwise correlation of principal components (PCs), or from the total length of the genome shared identical-by-descent (IBD) between pairs of individuals. PCA and GRM can be further computed from different sets of genetic variants (e.g. common or rare).

##### Identity by descent inference

Pairwise long IBD (>5cM) sharing was estimated from released phased data using Germline2^46^ using autosomal biallelic SNPs with MAF > 0.01 for IBD estimation. We removed variants with more than 10% missing data and samples with more than 10% missingness. Then, for each pair of individuals, we computed the total length of the shared segments between two individuals as the input for network construction and community detection. We removed related individuals using the list provided in the metadata from the jointly called.

##### GRM estimation

The Genetic Relationship Matrix (GRM) was estimated using GCTA (v1.94.1)^47^ using autosomal biallelic SNPs. We removed variants with more than 10% of missing data and those failing the Hardy-Weinberg equilibrium test (p-value < 1e-10). We also removed samples with more than 10% missingness (No samples were removed). We pruned variants for linkage disequilibrium in Plink (v1.90b6.21)^48^ (with --indep-pairwise 50, 5, 0.2). We estimated GRM matrices separately from common (MAF > 1%), and rare (MAF < 1%) variants, excluding singletons, referring to them as common- and rare-GRM, respectively.

##### PCA correlation

The PCs were made available as part of the metadata in the joint 1KGP + HGDP variant call set (https://gnomad.broadinstitute.org/help/hgdp-1kg-annotations). We used the first 20 PCs to compute pairwise genetic similarity (Pearson correlation) between individuals, setting negative correlations to zero.

Our pipeline outputs a graphic representation of the built network with different features. We used the Fruchterman-Reingold layer to aid in the visualization of dense data points in the network^49^. This algorithm emulates a particle system, where the vertices represent charged particles that repel each other, while the edges represent springs that attract the connected vertices. Through multiple iterations, the algorithm fine-tunes and provides the positions of the vertices to attain a state of equilibrium^49^.

### Louvain Algorithm for Community Detection

For each genetic metric, we used the Louvain algorithm^31^ for community detection. The algorithm partitions a network into communities, or modules, which are groups of nodes that are more densely connected than would be expected by chance. This algorithm employs a two-phase iterative approach to determine the community structure that maximizes modularity, which measures the level of connectivity within these communities. In the initial iteration, each individual is considered a community. Then, during phase one, it evaluates whether moving individuals from one community to another improves modularity. In phase two, it constructs a new network where the communities identified in phase one are treated as individuals. These phases are repeated until the modularity cannot be further improved.

We implemented the algorithm using the igraph package in R^45^. In this implementation modularity^50^ is defined as:

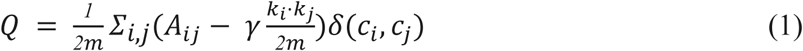

Where *m* is the weight of all network links, *A_ij_* is the sum of the weights of links that connect the node *i* with the node *j*, *k_i_* is the sum of the weights of links in node *i*, *k_j_* is the sum of the weights of links in node *j*, *Σ_i,j_* is the sum of the weights for all pairs of nodes *i* and *j*.

In this equation, *A_ij_* reflects the density of interactions between the pair of nodes *i* and *j*, and 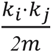 is the expected density by chance. Thus, the *γ* parameter determines the density threshold for nodes to be reassigned to communities identified by the algorithm. A smaller gamma yields a small number of larger communities due to many nodes exceeding the density threshold. In contrast, a higher gamma leads to more, but smaller in size, communities, as only the denser nodes can surpass the density threshold. When the gamma parameter equals 1, the equation transforms into the standard equation for modularity.

The Louvain algorithm optimizes modularity, but also suffers from the resolution limit, making it challenging to detect smaller communities within the network^33,51^. To properly address these issues, we also implemented the Leiden algorithm in the GG-NC pipeline, which is not affected by the resolution limit (Figures S3 and S20)^30^. Louvain can also find poorly connected communities, and in the worst-case scenario, communities could be internally disconnected^33^. The Leiden algorithm overcomes this limitation by adding an extra step (refinement of the partition) to guarantee internally connected communities.

### *Resolution plot* based on community detection at multiple resolutions

We applied the Louvian community detection algorithm (described in more detail above) – a heuristic method that is based on modularity optimization^31^. We defined the exploration space of this parameter as a logarithmic space from −2 to 2 considering 50 steps. We refer to log_10(gamma) as the *resolution value*. The membership of the individuals to the emergent communities at each resolution value can be represented in a ‘*resolution plot*’ (Figure 2), which shows how individuals change their membership across the range of resolution values. Such a visualization is inspired from its prior use to visualize protein-protein interaction networks^32^. It is important to note that the nomenclature of the communities is maintained across resolution values and nodes are reordered on the x-axis to try to maintain the continuity of the communities as much as possible, using a convention for labeling communities described in Lewis et al (2010)^32^. For example, community 4 will be labeled and colored the same across resolutions, also individuals belonging to this community will be ordered together on the x-axis. Communities that do not have more than 6 members in any resolution are colored in white in the *Resolution Plot* (the smallest cohort we analyzed has 6 individuals).

### Assess community stability at each resolution and compare with super population structure

The pipeline can compute two measures of ‘stability’, which describes the extent to which individual memberships in communities are stable for a given resolution value. To do so we ran the Louvain algorithm 100 times for each resolution value and compared the communities obtained pairwise. We used the Adjusted Rand Index (ARI) and the Normalized Information Distance (NID) metrics. Additionally, we compared the 100 runs for each resolution against the super populations using the same metrics.

We implemented the functions NID() and ARI() in the aricode R package, both highly efficient for their respective purposes. However, specific considerations arise in trivial cases that require attention:

For NID(), when each individual in both partitions form their own community, the output is “0”. When all individuals in both partitions belong to a single community, the result is “NaN”. For ARI(), when each individual in both objects forms their own community, the function produces “NaN”. When all individuals in both partitions belong to a single community, the output is “1”.

#### Normalized information distance (NID)

To evaluate the stability of community formation using the Louvain Algorithm method, we employed the Normalized Information Distance (NID)^52^, as a measure to quantify the resemblance in the distribution of individuals across communities, using the function NID() in the aricode R package^53^. This measure, based on information entropy, was calculated based on 100 iterations of the algorithm for each resolution value.

The general formula for the NID between two objects *X* and *Y* is expressed as:

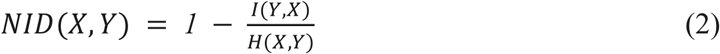

Where *H* is entropy: *H*(*X*) = −*Σ_i_p_i_*(*X*)*logp_i_*(*X*). Thus, the mutual information is *I*(*Y*, *X*) = −*H*(*X*|*Y*) + *H*(*X*) + *H*(*Y*) and *H*(*X*, *Y*) is the joint entropy. The function is normalized to fit the range [0, 1], where 0 means that the two objects are identical and 1 that they are completely different.

#### Adjusted Rand Index (ARI)

We also used the Adjusted Rand Index (ARI), an extension of the Rand Index^54^ as a second external cluster validation. The Rand Index (RI) was created by Rand in 1971 as a measure to evaluate the similarity between clustering and classifications.

Considering two objects *X* = {*X_1_*, *X_2_*, … *X_n_*} and *Y* = {*Y_1_*, *Y_2_*, … *Y_n_*}, we can build a contingency matrix *M* where every column represents an element of *X*, every row represents an element of *Y*, *n* is the length of the objects, and the entries *m_ij_* indicate the overlap between X and Y. Then, *m_i._* represents the sum over the ith row, *m*_.*j*_ is the sum over the jth column. The equation for ARI estimation is given by:

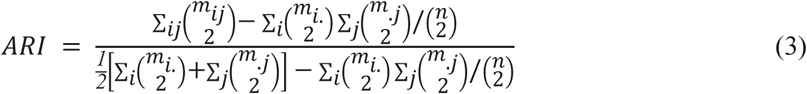

The ARI value ranges from −1 to 1, where a score of 1 denotes a perfect match between the partitions, 0 indicates expected similarity by chance, and −1 perfect disagreement.

Besides using these metrics, we also analyzed the results for a given resolution value, summarizing the 100 runs in a single heatmap. The heatmap represents a squared matrix in which columns and rows are the individuals. The value indicates how many times a pair of individuals were grouped in the same community. Thus a value of 100 means that the pair of individuals were always in the same community. The heatmaps were generated using ComplexHeatmap^55^ (2.14.0) library from R.

### Wilcoxon test

We performed a no-paired one-side Wilcoxon rank-sum test using the R function wilcox.test() (alternative = “greater”) to determine whether the distribution of ARI values was significantly greater for (1) between communities, which quantifies the consistency of individual membership across runs at each resolution, than (2) communities vs super populations, which measures the similarity between the communities and the predefined super populations.

### Community networks for a given resolution

We built three-dimensional (3D) community networks by calculating the average x, y, and z coordinates across individuals within each community. To do this, we first utilized the Fruchterman-Reingold algorithm to determine the 3D layout of individuals and computed the x, y, and z coordinates for each community by averaging across the coordinates of individuals in that community leveraging the ucie package in R^56^. We assigned colors to communities based on two different methodologies (see below).

### Color coding

We employed two methodologies for assigning colors to communities that use CIELab color space, a three-dimensional color model aimed at accurately representing the diverse range of colors observed by the human eye in a consistent and unbiased manner. The first method allowed us to obtain distinct colors that clearly differentiate each community in a network using the distinct_colors() function from the chameleon package. However, is it possible for colors to provide information about the genetic closeness of communities? For the second methodology, we first explored different resolutions to identify the one where all communities are present simultaneously. Then, we leveraged the Fruchterman-Reingold algorithm’s capability to assign 3D relative positions based on community connections, and we used the data2cielab() function from the ucie package that retrieves the corresponding color for each community based on its placement in a three-dimensional space. Communities are checked for any omissions, as the highest resolution may not encompass all. They are then aggregated by averaging their positions across the resolutions where they appear. The latter method enabled us to observe genetically close communities with colors that are more similar, and vice versa (Figures S5 and S6).

### Visualizing communities as resolution changes

Finally, we developed a shiny app to make our results more interactive and allow engagement with scientists and thee general public alike. The shiny app allows us to see the different communities that emerge at different resolution values and their geographic distribution across the analyzed genetic similarity metrics (IBD, PCA, GRM common and GRM rare). Each pie chart shows the proportion of the individuals that belong to each community. A slider allows users to try different resolution values while displaying the number of detected communities and their community network composition. We also offer the option to change the *resolution plot* and map colors, so that similar colors indicate closeness between communities.

Our browser has a “Customize” panel where users can upload the output files generated with our GG-NC pipeline to analyze and visualize results on their own genetic datasets. Since our goal is to make a user-friendly app, we also offer a video tutorial in two languages (English and Spanish) that explains and exemplifies the applications of our browser.

### Community detection on variants associated with particular traits and diseases

Using autosomal biallelic SNPs, we removed variants with more than 10% missingness. We also removed samples with more than 10% of missingness and related individuals. We kept variants inside the genomic coordinates of genes associated with the following traits:

- **Altitude (7 genes):** EPAS1, EGLN1, PPARA, CBARA1, VAV3, ARNT2, and *THRB*^38^.
- **Type 2 diabetes (15 genes):** HNF4A, RREB1, GCKR, POC5, ANKH, WSCD2, KCNJ11, PAM, TM6SF2, LPL, PLCB3, SLC30A8, PNPLA3, HNF1A, and *GIPR*^36^.
- **Skin pigmentation (38 genes):** OCA2, SLC24A5, SLC45A2, TYR, MFSD12, DDB1, TMEM138, HERC2, IRF4, BEND7, PRPF18, MC1R, ASIP, TYRP1, SMARCA2, VLDLR, SNX13, GRM6, ATF1, WNT1, SILV, OPRM1, EGFR, ZNF804B, PDE4B, RIPK5, PA2G4P4, PPARGC1B, AHR, AGR3, TRPS1, BNC2, EMX2, TPCN2, DCT, ATP11A, SLC24A4, and *KIAA0930*^37^.

We pruned variants for linkage disequilibrium in Plink (v1.90b6.21) (with --indep-pairwise 50, 5, 0.1). The first 20 PCs were estimated for each subset of variants using smartpca (v13050) from Eigensoft (v6.0.1)^57,58^ (using numoutlieriter: 5, numoutlierevec: 10, outliersigmathresh: 6, and qtmode: 0).

### IBD network modifications

Aiming to improve the visualization of the networks shown in this paper we modified the IBD network generated, iteratively removing individuals (nodes) that were only connected to a single node or were completely disconnected. Further, we removed individuals that were isolated from the overall network (forming communities of <25 members even at the low resolution of R=-2)(Supplementary Tables 2 and 3). The results of the network without outlier removal can be seen on our web browser. In the manuscript, we present stability results on the IBD network after outlier removal (Figure 3); however, stability results are qualitatively the same on the full network (Figure S12).

### ADMIXTURE and PCA analysis

For consistency, only individuals included in the final IBD network were considered for ADMIXTURE and PCA analyses (Figure 4). We removed variants with more than 10% missingness and MAF < 0.05. We pruned variants for linkage disequilibrium in Plink (v1.90b6.21) (with --indep-pairwise 100, 10, 0.1).

ADMIXTURE (V1.3.0) was run from K=5 to K=25 estimating the cross-validation error. Results were plotted using pong (v1.5)^59^. The first 20 PCs were estimated using smartpca (v13050) from Eigensoft (v6.0.1). Results were plotted using R.

## Acknowledgments

This project was supported by CONAHCYT Ciencia de Frontera (Frontiers of Science) project no. 319349 in the modality “Paradigmas y Controversias de la Ciencia 2022”. M.J.P-M. is a doctoral student from Programa de Doctorado en Ciencias Biomédicas, Universidad Nacional Autónoma de México (UNAM) and received a fellowship (889218) from CONAHCYT. Y.S.P-G. and C. Q. L. were supported by CONAHCYT project no. 319349. A.A.Z. was supported by the National Institute of General Medical Sciences award R00GM137076. A.C.F.L. was supported by 1K99HG012809 award from NHGRI. We also want to thank Brandon Ezequiel Bello Chimal for his support in the recording of the tutorial videos for the GG-NC browser.

## Author Contributions

M.S. conceived the project. M.J.P.M, Y.P.G, B.E.L.A and C.Q.L performed analyses, created the computational pipeline and the web browser. A.C.F.L, K.A.B, T.L, A.Z. and M.S. provided conceptual and technical input throughout the project. All authors wrote and edited the paper.

## Data Availability

1000 Genomes and Human Genome Diversity Project data analyzed in this dataset was downloaded from gnomAD (https://gnomad.broadinstitute.org/downloads#v3-hgdp-1kg). The result of our GG-NC pipeline on this dataset can be accessed on our web browser (https://sohail-lab.shinyapps.io/GG-NC/).

## Code Availability

The GG-NC pipeline is available through our GitHub repository (https://github.com/mariajpalma/GG-NC). The web browser for the Global Genome Network Communities is available at https://sohail-lab.shinyapps.io/GG-NC/ and can be used for further exploration of our results, as well as to visualize results for any genetic dataset that can be analyzed using our GitHub repository.

## Extended Data Table Captions

**Extended Data Table 1. Genes and coordinates for trait-specific analysis of Type 2 diabetes.** Genomic coordinates (GRch38) of genes associated with Type 2 diabetes^36^.

**Extended Data Table 2. Genes and coordinates for trait-specific analysis of skin pigmentation.** Genomic coordinates (GRch38) of genes associated with skin pigmentation^37^.

**Extended Data Table 3. Genes and coordinates for trait-specific analysis of high altitude adaptation.** Genomic coordinates (GRch38) of genes associated with adaptation to high altitude^38^.

## References

1. Rosenberg, N. A. et al. Genetic structure of human populations. Science 298, 2381–2385 (2002).

2. 1000 Genomes Project Consortium et al. A global reference for human genetic variation. Nature 526, 68–74 (2015).

3. Bird, K. A. & Carlson, J. Typological thinking in human genomics research contributes to the production and prominence of scientific racism. Front. Genet. 15, 1345631 (2024).

4. Terrell, J., Golitko, M., Dawson, H. & Kissel, M. Modeling the Past: Archaeology, History, and Dynamic Networks. (Berghahn Books, 2023).

5. Koenig, Z. et al. A harmonized public resource of deeply sequenced diverse human genomes. Genome Res. 34, 796–809 (2024).

6. Dunn, T. & Dobzhansky, T. Heredity, Race and Society. (Pelican Books, 1946).

7. Saini, A. Superior: The Return of Race Science. (Beacon Press, 2019).

8. Yudell, M. Race Unmasked: Biology and Race in the Twentieth Century. (Columbia University Press, 2014).

9. Bergström, A. et al. Insights into human genetic variation and population history from 929 diverse genomes. Science 367, eaay5012 (2020).

10. Mallick, S. et al. The Simons Genome Diversity Project: 300 genomes from 142 diverse populations. Nature 538, 201–206 (2016).

11. GenomeAsia100K Consortium. The GenomeAsia 100K Project enables genetic discoveries across Asia. Nature 576, 106–111 (2019).

12. Lewis, A. C. F. et al. An Ethical Framework for Research Using Genetic Ancestry. Perspect. Biol. Med. 66, 225–248 (2023).

13. National Academies of Sciences, Engineering, and Medicine; Division of Behavioral and Social Sciences and Education; Health and Medicine Division; Committee on Population; Board on Health Sciences Policy; Committee on the Use of Race, Ethnicity, and Ancestry as Population Descriptors in Genomics Research. Using Population Descriptors in Genetics and Genomics Research: A New Framework for an Evolving Field. (National Academies Press (US), Washington (DC), 2023).

14. Panofsky, A., Dasgupta, K. & Iturriaga, N. How White nationalists mobilize genetics: From genetic ancestry and human biodiversity to counterscience and metapolitics. Am. J. Phys. Anthropol. 175, 387–398 (2021).

15. Jedidiah, B. M., Henn, D. R. & Al-Hindi, S. Counter the Weaponization of Genetics Research by Extremists. Nature 610, 444–447 (2022).

16. Wills, M. Are Clusters Races? A Discussion of the Rhetorical Appropriation of Rosenberg et al.’s ‘Genetic Structure of Human Populations’. Philos. Theory Pr. Biol. 9, (2017).

17. Alexander, D. H., Novembre, J. & Lange, K. Fast model-based estimation of ancestry in unrelated individuals. Genome Res. 19, 1655–1664 (2009).

18. Pritchard, J. K., Stephens, M. & Donnelly, P. Inference of population structure using multilocus genotype data. Genetics 155, 945–959 (2000).

19. Terrell, J. E. Social network analysis of the genetic structure of Pacific islanders. Ann. Hum. Genet. 74, 211–232 (2010).

20. Marcus, J. H. & Novembre, J. Visualizing the geography of genetic variants. Bioinformatics 33, 594–595 (2016).

21. Biddanda, A., Rice, D. P. & Novembre, J. A variant-centric perspective on geographic patterns of human allele frequency variation. Elife 9, e60107 (2020).

22. Diaz-Papkovich, A., et al. Topological stratification of continuous genetic variation in large biobanks. bioRxiv (2023) doi:10.1101/2023.07.06.548007.

23. Grundler, M. C., Terhorst, J. & Bradburd, G. S. A geographic history of human genetic ancestry. bioRxiv (2024) doi:10.1101/2024.03.27.586858.

24. Donovan, B. M. et al. Toward a more humane genetics education: Learning about the social and quantitative complexities of human genetic variation research could reduce racial bias in adolescent and adult populations. Sci. Educ. 103, 529–560 (2019).

25. Greenbaum, G., Rubin, A., Templeton, A. R. & Rosenberg, N. A. Network-based hierarchical population structure analysis for large genomic data sets. Genome Res. 29, 2020–2033 (2019).

26. Greenbaum, G., Templeton, A. R. & Bar-David, S. Inference and Analysis of Population Structure Using Genetic Data and Network Theory. Genetics 202, 1299–1312 (2016).

27. Belbin, G. M. et al. Toward a fine-scale population health monitoring system. Cell 184, 2068–2083.e11 (2021).

28. Kuismin, M. O., Ahlinder, J. & Sillanpӓӓ, M. J. CONE: Community Oriented Network Estimation Is a Versatile Framework for Inferring Population Structure in Large-Scale Sequencing Data. G3 Genes|Genomes|Genetics 7, 3359–3377 (2017).

29. Koyama, S. et al. Decoding genetics, ancestry, and geospatial context for precision health. medRxiv (2023) doi:10.1101/2023.10.24.23297096.

30. Traag, V. A., Van Dooren, P. & Nesterov, Y. Narrow scope for resolution-limit-free community detection. Phys Rev E Stat Nonlin Soft Matter Phys 84, 016114 (2011).

31. Blondel, V. D., Guillaume, J.-L., Lambiotte, R. & Lefebvre, E. Fast unfolding of communities in large networks. J. Stat. Mech. 2008, P10008 (2008).

32. Lewis, A. C. F., Jones, N. S., Porter, M. A. & Deane, C. M. The function of communities in protein interaction networks at multiple scales. BMC Syst. Biol. 4, 100 (2010).

33. Traag, V. A., Waltman, L. & van Eck, N. J. From Louvain to Leiden: guaranteeing well-connected communities. Sci Rep 9, 5233 (2019).

34. Palamara, P. F., Lencz, T., Darvasi, A. & Pe’er, I. Length distributions of identity by descent reveal fine-scale demographic history. Am. J. Hum. Genet. 91, 1150 (2012).

35. Zaidi, A. A. & Mathieson, I. Demographic history mediates the effect of stratification on polygenic scores. Elife 9, e61548 (2020).

36. Mahajan, A. et al. Refining the accuracy of validated target identification through coding variant fine-mapping in type 2 diabetes. Nat. Genet. 50, 559–571 (2018).

37. Quillen, E. E. et al. Shades of complexity: New perspectives on the evolution and genetic architecture of human skin. Am. J. Phys. Anthropol. 168 Suppl 67, 4–26 (2019).

38. Scheinfeldt, L. B. et al. Genetic adaptation to high altitude in the Ethiopian highlands. Genome Biol. 13, 1–9 (2012).

39. He, G. et al. A comprehensive exploration of the genetic legacy and forensic features of Afghanistan and Pakistan Mongolian-descent Hazara. Forensic Sci. Int. Genet. 42, e1–e12 (2019).

40. Mohsen, H. et al. Dynamic clustering of genomics cohorts beyond race, ethnicity--and ancestry. (2023) doi:10.1101/2023.08.04.552035.

41. Jablonski, N. G. Skin color and race. Am J Phys Anthropol 175, 437–447 (2021).

42. Kaplan, J. & Winther, R. Realism, Antirealism, and Conventionalism about Race. Philos. Sci. (Paris) 81, 1039–1052 (2014).

43. Ávila-Arcos, M. C. et al. Population History and Gene Divergence in Native Mexicans Inferred from 76 Human Exomes. Mol. Biol. Evol. 37, 994–1006 (2020).

44. Caggiano, C. et al. Disease risk and healthcare utilization among ancestrally diverse groups in the Los Angeles region. Nat. Med. 29, 1845–1856 (2023).

## Additional References for Materials and Methods

45. Csardi, G. & Nepusz, T. The igraph software. Complex syst 1695, 1–9 (2006).

46. Nait Saada, J., et al. Identity-by-descent detection across 487,409 British samples reveals fine scale population structure and ultra-rare variant associations. Nat. Commun. 11, 6130 (2020).

47. Yang, J., Lee, S. H., Goddard, M. E. & Visscher, P. M. GCTA: a tool for genome-wide complex trait analysis. Am. J. Hum. Genet. 88, 76–82 (2011).

48. Chang, C. C. et al. Second-generation PLINK: rising to the challenge of larger and richer datasets. Gigascience 4, 7 (2015).

49. Fruchterman, T. M. J. & Reingold, E. M. Graph drawing by force-directed placement. Softw. Pract. Exp. 21, 1129–1164 (1991).

50. Clauset, A., Newman, M. E. J. & Moore, C. Finding community structure in very large networks. Phys. Rev. E Stat. Nonlin. Soft Matter Phys. 70, 066111 (2004).

51. Fortunato, S. & Barthélemy, M. Resolution limit in community detection. Proceedings of the National Academy of Sciences 104, 36–41 (2007).

52. Kraskov, A., Stögbauer, H., Andrzejak, R. G. & Grassberger, P. Hierarchical clustering using mutual information. EPL 70, 278–284 (2005).

53. Chiquet, J., Rigaill, G. & Sundqvist, M. Aricode: Efficient computations of standard clustering comparison measures. CRAN: Contributed Packages The R Foundation 10.32614/cran.package.aricode (2018).

54. Hubert, L. & Arabie, P. Comparing partitions. J. Classif. 2, 193–218 (1985).

55. Gu, Z. Complex heatmap visualization. Imeta 1, e43 (2022).

56. Koutrouli, M., Morris, J. H. & Jensen, L. J. U-CIE [/ju: ’si:/]: Color encoding of high-dimensional data. Protein Sci 31, e4388 (2022).

57. Patterson, N., Price, A. L. & Reich, D. Population Structure and Eigenanalysis. PLoS Genet. 2, e190 (2006).

58. Price, A. L. et al. Principal components analysis corrects for stratification in genome-wide association studies. Nat. Genet. 38, 904–909 (2006).

59. Behr, A. A., Liu, K. Z., Liu-Fang, G., Nakka, P. & Ramachandran, S. pong: fast analysis and visualization of latent clusters in population genetic data. Bioinformatics 32, 2817–2823 (2016).

