## Supplementary material for "The multi-scale complexity of human genetic variation beyond continental groups"

### Table of Contents

**List of Supplementary Figures** **3**
**List of Supplementary Tables** **5**
**Supplementary text** **6**
Stability of communities 6
Community detection using trait-associated genetic variation 6
**Supplementary Figures** **9**
**Supplementary Table Captions** **39**

### List of Supplementary Figures

**Figure S1. Pipeline workflow.**
**Figure S2. Graphical description of the quality control and estimation of genetic metrics.**
**Figure S3. *Resolution plot* for IBD results and associated individual and community networks, along with the geographic distribution of the communities using Leiden**
**Figure S4. Differences in the visualization of the network of individuals excluding and including individuals from smaller communities.**
**Figure S5. *Resolution plot* for IBD networks and geographic distribution of the communities using similar colors.**
**Figure S6. Community networks based on IBD using similar colors**.
**Figure S7. Individual networks colored by community based on PCA and GRM common and rare variants.**
**Figure S8. Community networks based on PCA and GRM common and rare variants.**
**Figure S9. Communities detected in the IBD-network are fairly stable across resolutions, and more stable at higher resolutions using Normalized Information Distance (NID).**
**Figure S10. Heatmap representation of the stability of the communities detected at different resolution values.**
**Figure S11. Geographic distribution of communities with the highest median coincidence with super populations (IBD, PCA, and GRM).**
**Figure S12. Stability analysis with Adjusted Rand Index (ARI) for communities detected in the GRM based on rare and common variants, and PCA networks at different resolutions.**
**Figure S13. Stability analysis with Normalized Information Distance (NID) for communities detected in the GRM based on rare and common variants, and PCA networks at different resolutions.**
**Figure S14. Admixture analysis using the merged dataset of the 1000 Genomes Project and HGDP at K=5 to 15.**
**Figure S15. Admixture analysis using the merged dataset of the 1000 Genomes Project and HGDP at K=16 to 25.**
**Figure S16. Coefficient of variation (CV) Error for different values of K in the ADMIXTURE analysis.**
**Figure S17. Overlap heatmap between communities at resolution value -2 and cohorts from 1000G and HGDP**.

**Figure S18. Overlap heatmap between communities at resolution value -0.041 and cohorts from 1000G and HGDP**.
**Figure S19. Components detected in the ADMIXTURE analysis at K=13 across the communities.**
**Figure S20. *Resolution plots* (Leiden algorithm) from networks using different definitions of genetic similarity and different subsets of genetic variants reveal different aspects of genetic relatedness**
**Figure S21. Geographic distribution of communities based on GRM or PCA.**
**Figure S22. Community formation in Hazara and Uygur Cohorts with increasing resolution in the GRM (common variants).**
**Figure S23. Community formation in Hazara and Uygur Cohorts with increasing resolution in the PCA.**
**Figure S24. Substructure in Africa detected on trait-specific results for the PCA networks.**
**Figure S25. The Americas in the trait-specific results for the PCA networks**.
**Figure S26. Substructure Variation Across Genomic Regions Associated with Different Traits.**
**Figure S27. Heatmaps of shared membership in trait-specific PCA communities.**
**Figure S28. Stability of communities for PCA results for Type 2 diabetes, Skin pigmentation, and High altitude at any given resolution is shown with Adjusted Rand Index (ARI).**
**Figure 29. Stability of communities for PCA results for Type 2 diabetes, Skin pigmentation, and High altitude at any given resolution is shown with Normalized Information Distance (NID).**
**Figure S30. Geographic distribution of communities at the resolution with the highest median coincidence with super populations (PCA trait specific).**

### List of Supplementary Tables

**Supplementary Table 1. IBD segments shared pairwise.**
**Supplementary Table 2. Sample metadata and inclusion on the different metrics.**
**Supplementary Table 3. Cohort sizes for the different metrics.**
**Supplementary Table 4. Wilcoxon results for community detection based on IBD.**
**Supplementary Table 5. Wilcoxon results for community detection based on PCA.**
**Supplementary Table 6. Wilcoxon results for community detection based on GRM common variants.**
**Supplementary Table 7. Wilcoxon results for community detection based on GRM rare variants.**
**Supplementary Table 8. Wilcoxon results for community detection based on trait-specific analysis (PCA skin pigmentation).**
**Supplementary Table 9. Wilcoxon results for community detection based on trait-specific analysis (PCA high altitude adaptation).**
**Supplementary Table 10. Wilcoxon results for community detection based on trait-specific analysis (PCA Type 2 diabetes).**

**Supplementary Tables 1-10 are presented in an additional excel file.**

### Supplementary text

#### Stability of communities

Another way (in addition to that presented in Figure 3) to illustrate the stability of communities is through a heatmap, which represents the proportion of runs across which two individuals are grouped in the same community at a given resolution (Figure S10). Color bars in the rows and columns allow us to compare the communities identified against continental groupings. In Figure S10, we observe that individuals from single continental groups are not grouped in a single community even at a resolution of -2, but instead that subsets of individuals within these groups have a higher tendency to be in the same community. Also, some communities integrate individuals from different continental groups. These results further highlight the dynamic nature of groupings of humans, which cannot be resolved by relying only on continental level groupings. For instance, we observe that some individuals from Africa are grouped together with individuals from Central South Asia just a few times, however, this group of individuals from Africa (Bantu in Kenya, Luhya in Webuye and half of Bantu individuals in South Africa) are always clustered together at a resolution of -2 and up to a resolution of -0.041. Similarly, at a resolution of -0.041 we observe the Maya individuals in Mexico consistently cluster together. However, they only occasionally merge (approximately 40 times) with other individuals from the Americas and Europe. All Maya individuals remain in the same community until resolution 1.429. In general, as shown by the heatmaps at R = -0.041, R=1.02, and R = 2, detected communities from GG-NC continue to mix up continental grouping labels further as resolution increases.

#### Community detection using trait-associated genetic variation

Not all genes/regions of the genome reflect the same evolutionary history and so the genetic similarity of individuals will not be identical for all loci. We highlight that the communities relevant for a gene or a given set of genes (related to a phenotype of interest) may be different from one set of genes to the next. The groupings most relevant for genetic epidemiology depend on the specific genetic variation and trait under consideration. To demonstrate this, we analyze sets of specific genes involved in Type 2 Diabetes, skin pigmentation, or altitude adaptation at diverse resolutions using PCA-based networks (Extended Data Tables 1-3). For each of these phenotypes, the first major division separates East Asian cohorts from the rest of the world rather than distinguishing Sub-Saharan Africans (including Afrodescendent individuals) from other groups like in the genome-wide PCA-based network. These cohorts are only distinguished at finer resolutions in the trait networks (PCA Skin, R=0.041, Altitude and T2D, R=-0.122). The subsequent community identified in all the trait-PCA-based networks consists of individuals from Central South Asia, including individuals sampled in other regions, but with ancestries from Central South Asia (Sri Lankan Tamil and Indian Telugu in the UK and Gujarati Indians in Houston, Texas, USA). However, in the T2D network, cohorts from East Asia are merged with some individuals from Indigenous groups of the Americas (Pima, Maya, Karitiana, Surui) and a fraction of cosmopolitan individuals from Latin America. Additionally, for the three trait-PCA-based networks, larger communities, primarily consisting of Sub-Saharan Africans and African Americans, persist at higher resolution values (Figure S24). In contrast to the diversity of communities found in this region, Indigenous individuals from the Americas and a subset of cosmopolitan individuals from Latin America remain in the same community until they eventually divide into much smaller communities (Figure S25).

As the resolution value increases we can observe a considerable number of communities emerging, showing substructure inside geographic regions that may be relevant for trait-specific epidemiology. These communities are not reflecting whole cohorts, but instead highlight finer scale substructure (Figure S26). The substructure is observed simultaneously in groups such as Europe, Central South Asia, and East Asia. This is in contrast to what we observe in the genome-based networks, where we do observe substructure but it is not as extensive.

The networks based on the different sets of associated genes for Type 2 Diabetes (T2D), Skin pigmentation, and adaptation to high-altitude environments show varying patterns and communities that are likely relevant for genetic epidemiology and evolutionary history alike for these different traits. For instance, the *resolution plot* for the skin pigmentation PCA network shows less substructure inside Europe, East Asia, the Middle East, and Central South Asia in comparison to the Type 2 Diabetes and altitude adaptation networks (Figure S26). This implies a more heterogeneous fine-scale genetic structure likely relevant for the latter traits (Diabetes and altitude adaptation) compared to the former (skin pigmentation).

Similar to Mohsen et al (2023)[40], we believe that this approach can allow users to explore the community structure relevant for genetic variation associated with their trait of interest, to help identify trait-specific variant clustering and epidemiology that may not relate to continental categories.

### Supplementary Figures

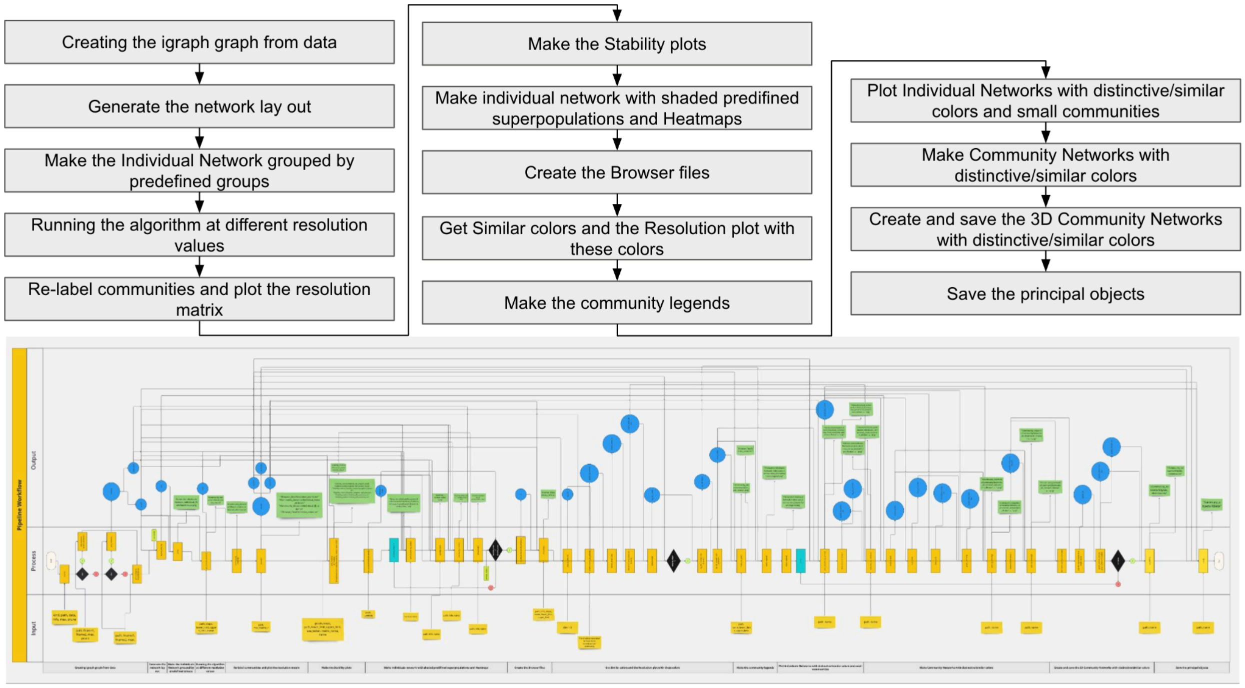


**Figure S1. Pipeline workflow.** This flowchart delineates the functions and operational framework of the pipeline used to generate various graphs, analyses, and visualizations. Inputs required for processing are represented as yellow rectangles on the left side, while the central section illustrates the functional processes, depicted as orange rectangles. On the right side, the output is represented by files shown in green rectangles and created objects indicated by blue circles. Arrows are used to indicate the connections among these components. Additional flowchart elements are present, including diamonds that signify decision points, along with their corresponding Yes/No options, and a counter displayed in a turquoise rectangle. The tabs on the left outline the different objectives of the pipeline. You can explore it in our GitHub repository.

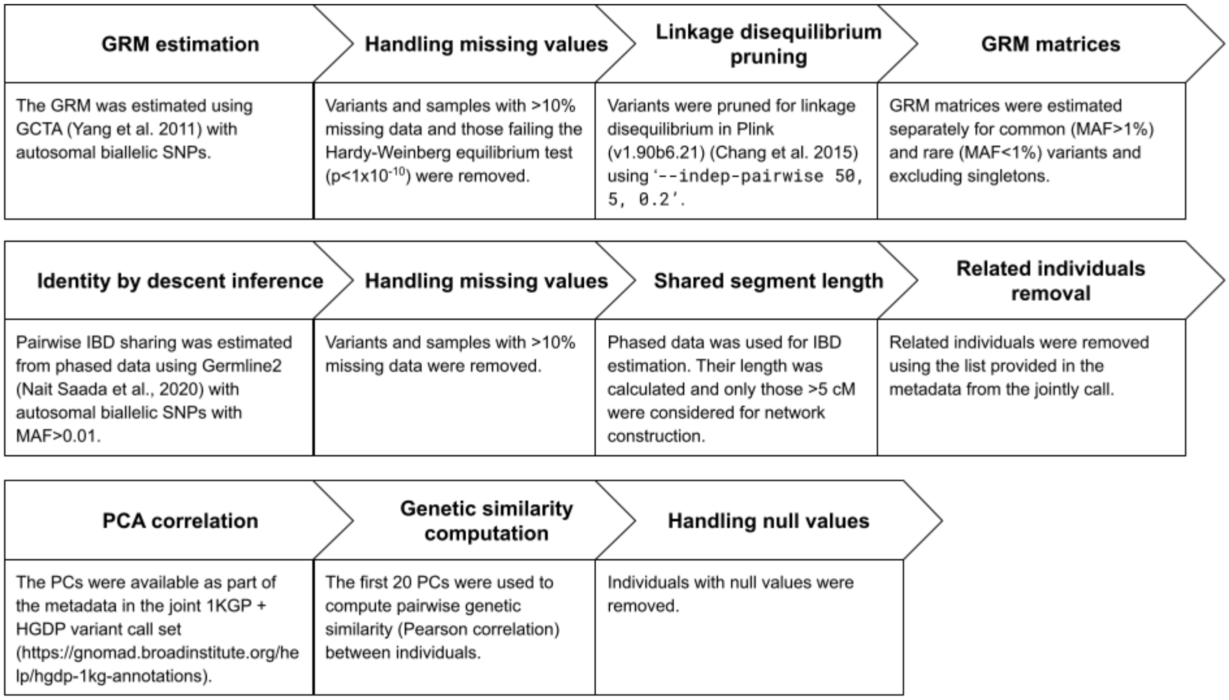


**Figure S2. Graphical description of the quality control and estimation of genetic metrics.** The chart illustrates the steps undertaken for the estimation and quality control of the Genetic Relationship Matrix (GRM), Identity by Descent (IBD), and Principal Component Analysis (PCA).

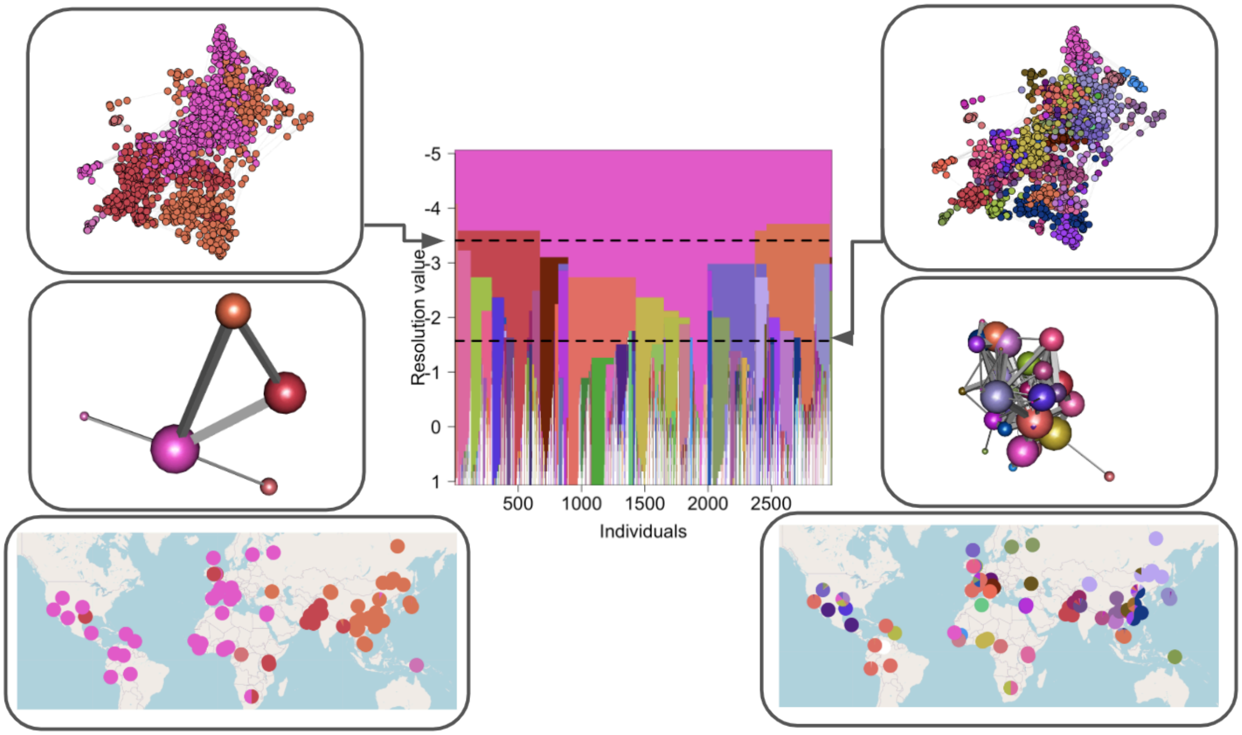


**Figure S3. *Resolution plot* for IBD results and associated individual and community networks, along with the geographic distribution of the communities using Leiden**. The central plot is the *Resolution plot* showing the results of the Leiden algorithm at 50 resolution values. The left panels display results for a resolution value of -3.4, and the right panel shows results at resolution -1.571. Each side includes (from top to bottom) the individual network, the community network, and the geographic distribution of the communities. Individual network is formed of 2,977 individuals represented by nodes (as for Louvain, 280 outlier samples were excluded from the network for visualization purposes, see methods and Figure S4) in which nodes are colored according to the community membership in the *resolution plot*. Community network plots present communities as nodes and the density of the connection among them as edges. In the maps, we show the 1000G project and HGDP cohorts using pie charts placed at sampling locations. Each pie chart represents the community membership of the individuals within each cohort.

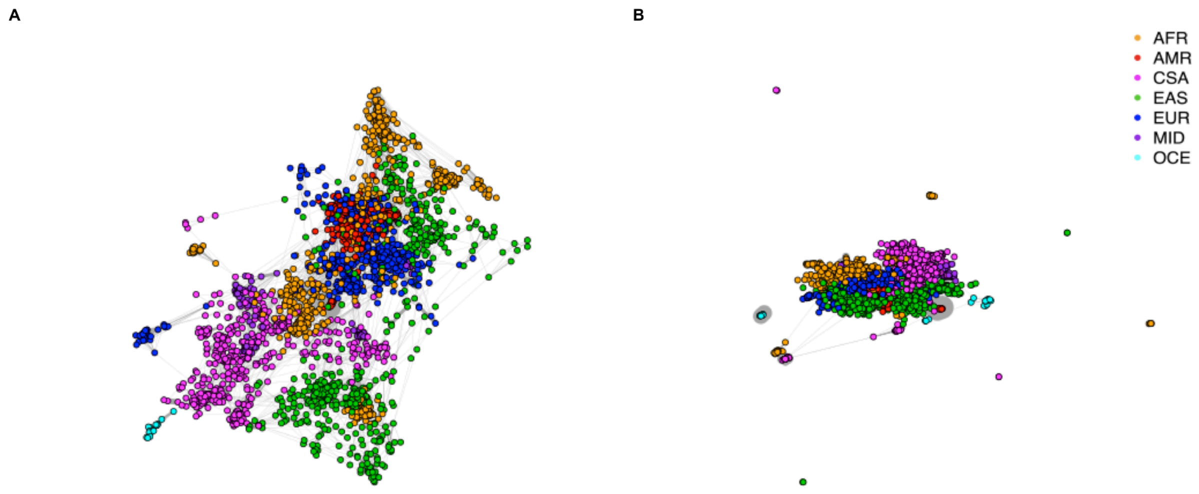


**Figure S4. Differences in the visualization of the network of individuals excluding and including individuals from smaller communities.** The IBD Individual networks are colored by super populations including the datasets from the 1000 Genomes Project and HGDP. **A)** Individual network excluding 280 samples for visualization purposes. In this case, nodes with a degree of 0 and 1 were excluded iteratively, along with communities containing fewer than 25 individuals, at a resolution of -2. **B)** Individual network including all the samples.

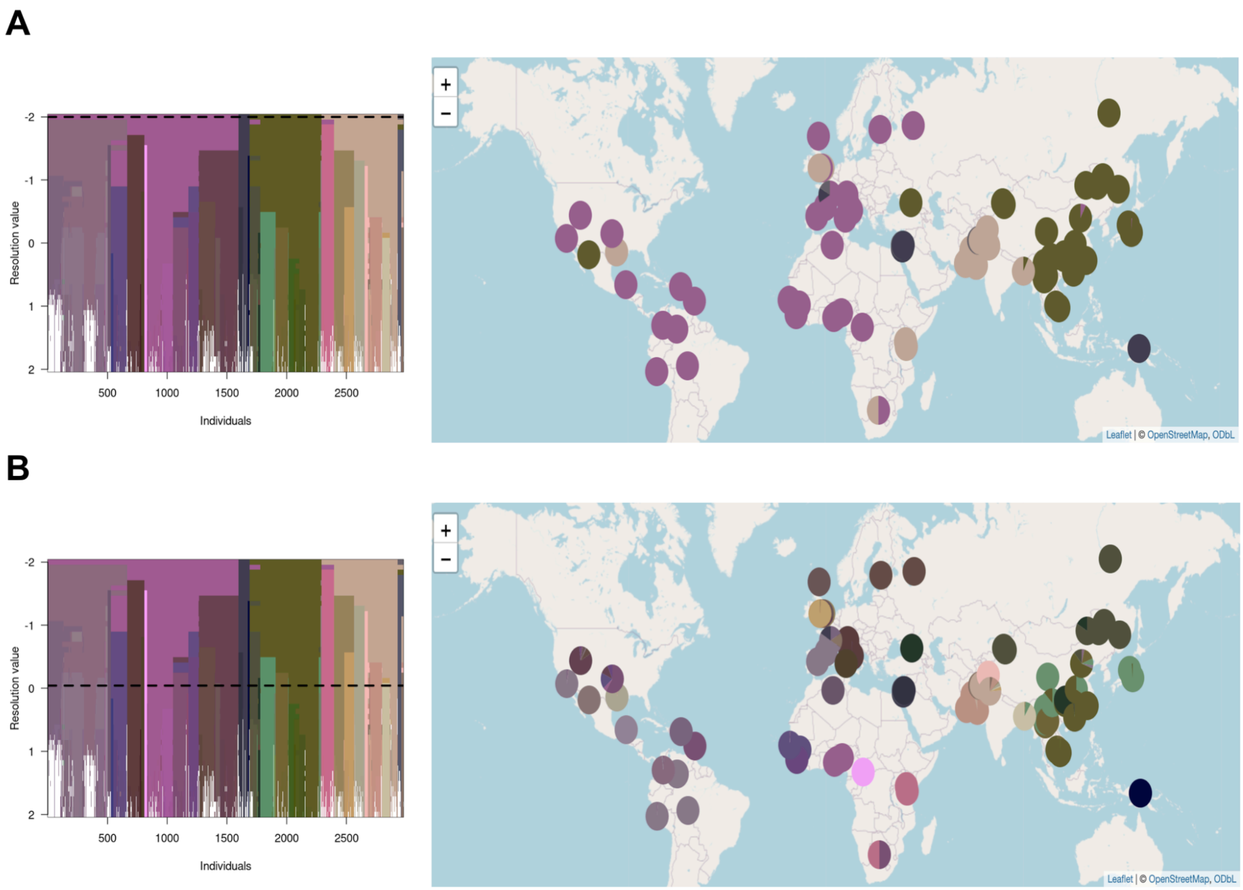


**Figure S5. *Resolution plot* for IBD networks and geographic distribution of the communities using similar colors.** The "similar colors" option allows coloring the communities similarly according to their proximity. **A)** *Resolution plot* and world map for resolution=-2. **B)** *Resolution plot* and world map based on IBD at a resolution of -0.041.

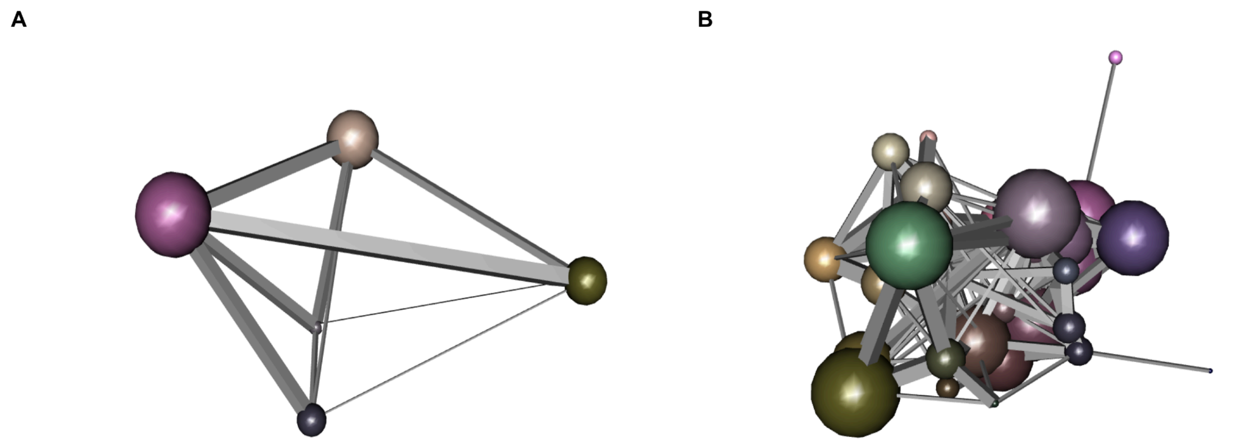


**Figure S6. Community networks based on IBD using similar colors**. **A)** Community network at a resolution of -2. **B)** Community network at a resolution of -0.41. Genetically closer communities are colored with a similar color.

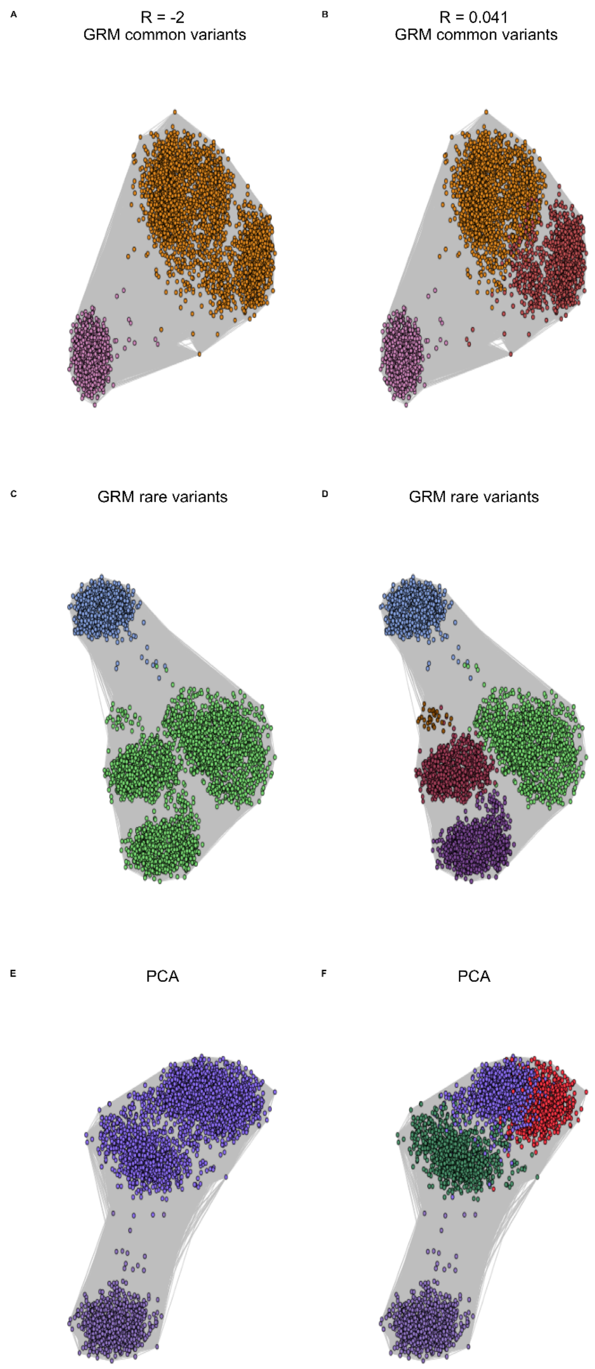


**Figure S7. Individual networks colored by community based on PCA and GRM common and rare variants. A,B)** Individual networks for the GRM (common variants) at a resolution of -2 and -0.041, respectively. **C,D)** Individual networks for the GRM (rare variants) at a resolution of -2 and -0.041, respectively. **E,F)** Individual networks for the PCA at a resolution of -2 and -0.41, respectively.

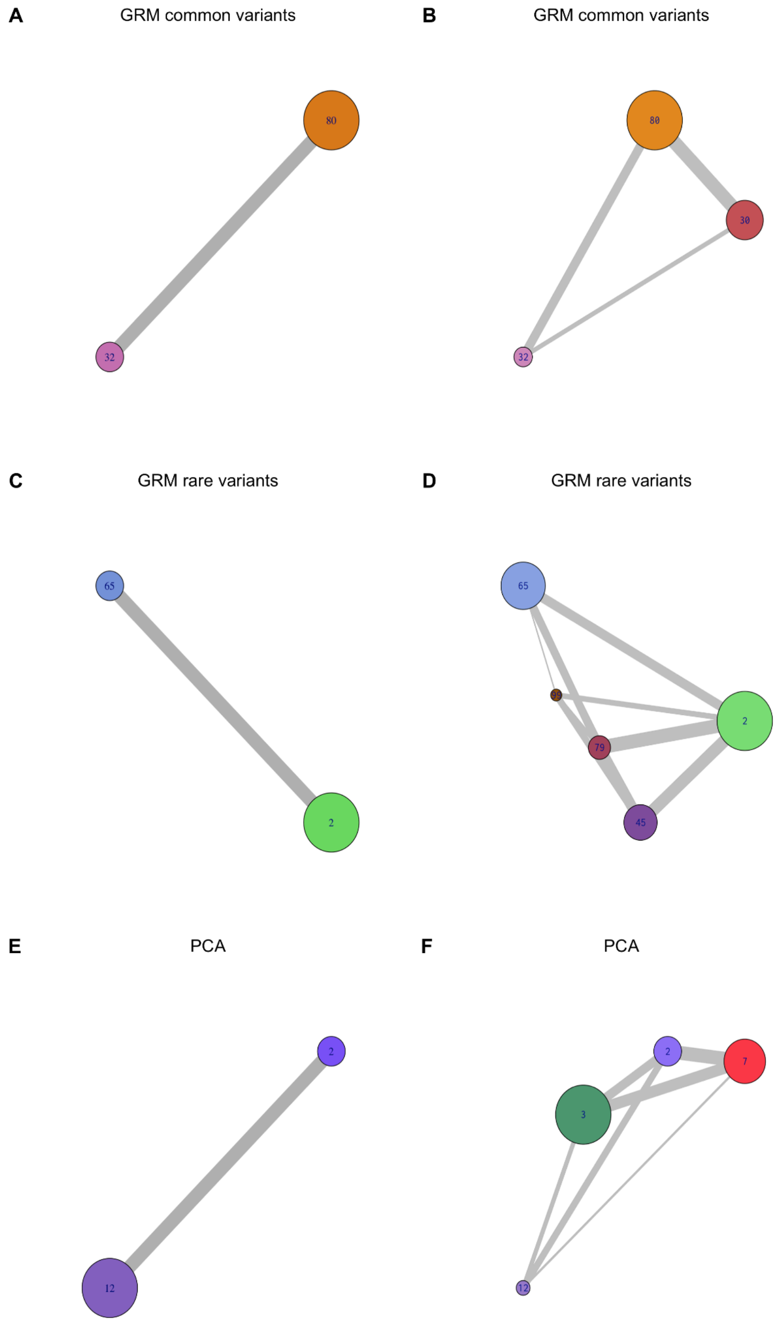


**Figure S8. Community networks based on PCA and GRM common and rare variants.** Community networks for the GRM (common variants) at resolutions of -2 and -0.041 (**A,B**); for the GRM (rare variants) at -2 and -0.041 (**C,D**); and for the PCA at -2 and -0.41 (**E,F**).

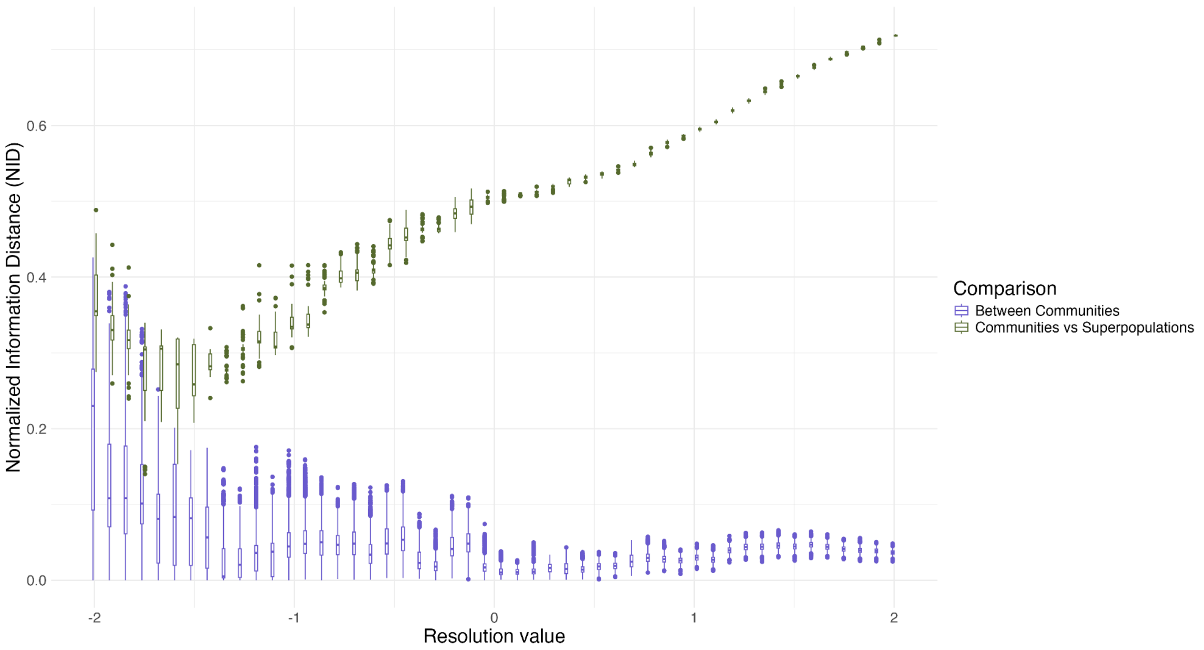


**Figure S9. Communities detected in the IBD-network are fairly stable across resolutions, and more stable at higher resolutions using Normalized Information Distance (NID).** The x-axis shows the resolution value. The y-axis shows the NID values. NID values closer to 0 indicate more individuals falling in the same communities across runs at a given resolution value. Purple boxplots summarize the comparison of community detection results across 100 independent runs at each resolution (see methods). Green boxplots represent the comparison between the independent runs and the super populations. In this case, NID values closer to 0 indicate greater similarity between the detected communities and the super populations. Boxplot elements: center line, median; box limits, upper and lower quartiles; whiskers, 1.58x interquartile range; points, outliers.

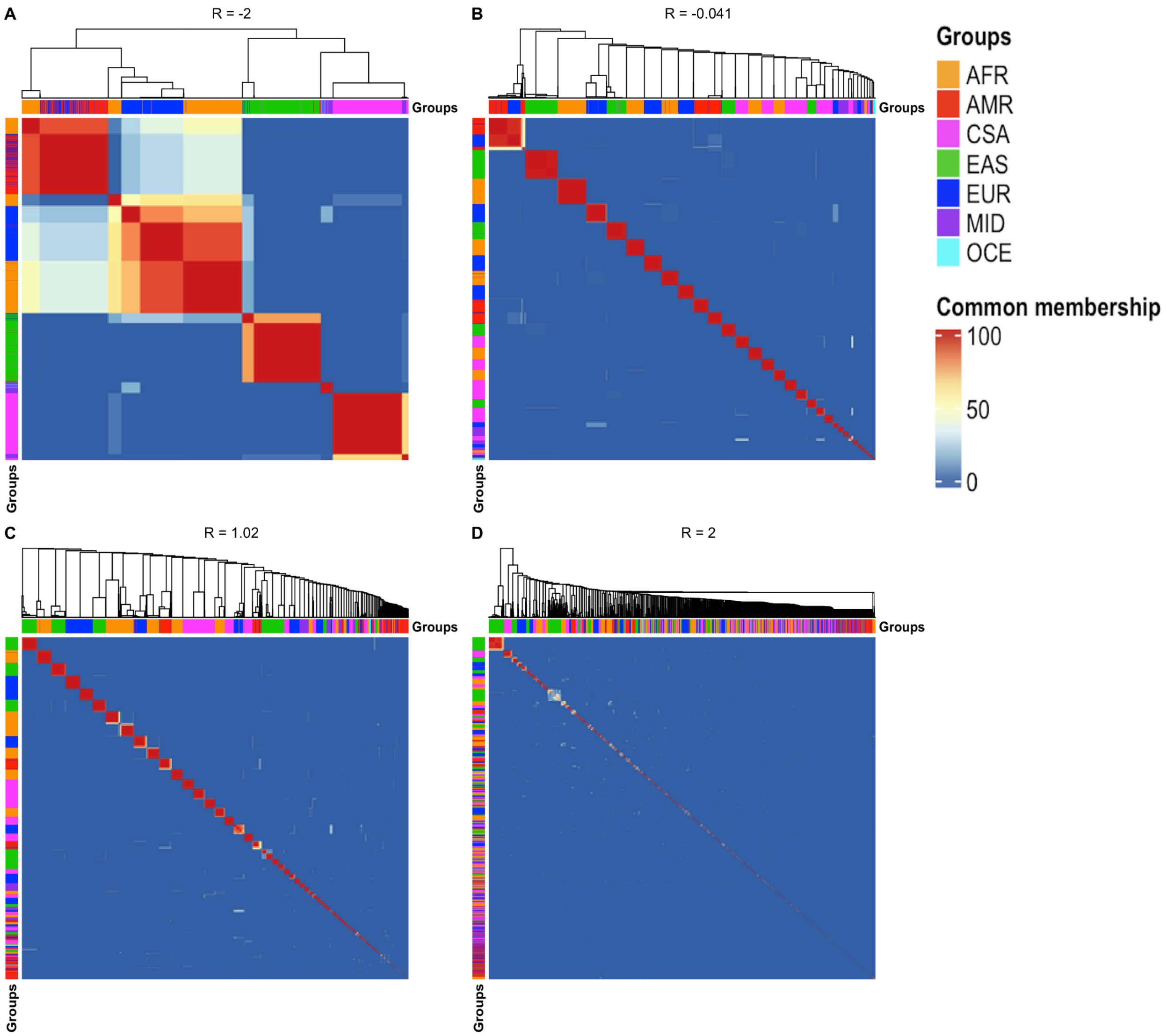


**Figure S10. Heatmap representation of the stability of the communities detected at different resolution values.** Heatmap representation of the stability of the communities detected at a resolution value -2, -0.041, 1.02, and 2. Each row and column corresponds to an individual. If individuals share the same community in the 100 runs, the intersection is shown in red, if they are always in different communities the intersection is shown in blue. The order of rows and columns is the same and is determined by the Euclidean distance. The annotation lines correspond to the super populations defined in the 1000 genomes project and HGDP.

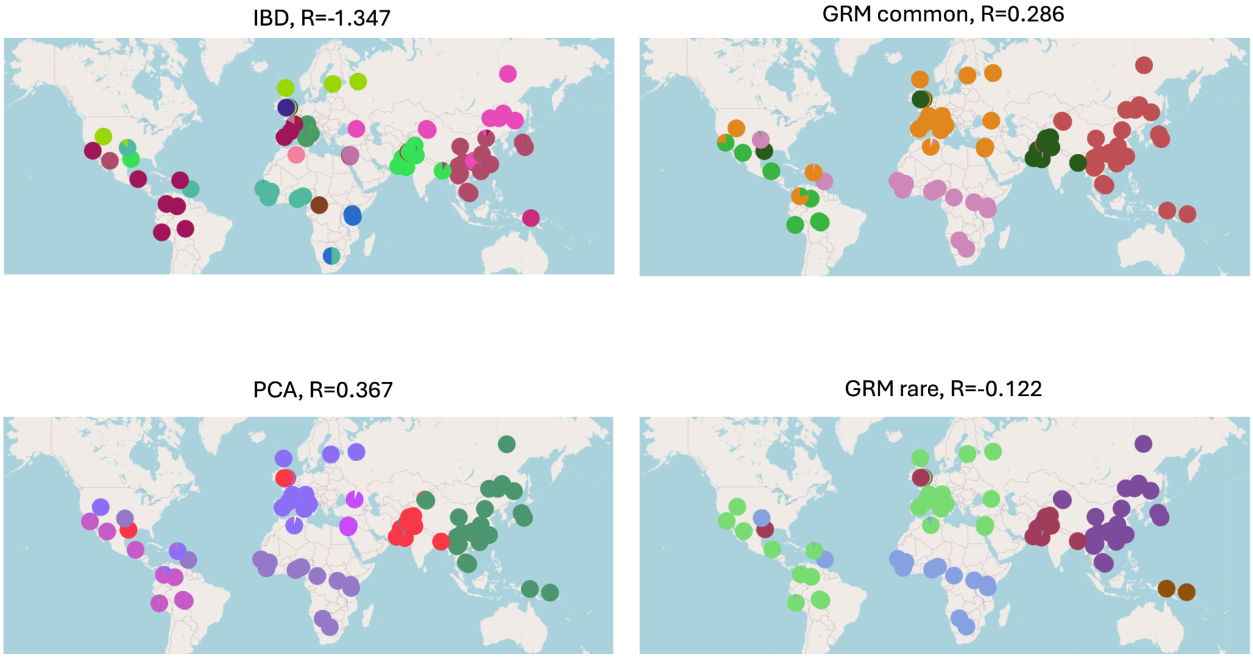


**Figure S11. Geographic distribution of communities with the highest median coincidence with super populations (IBD, PCA, and GRM).** Geographic distribution of detected communities for IBD, PCA, GRM common and rare variants at the resolution where the highest median ARI is observed when comparing communities with super populations.

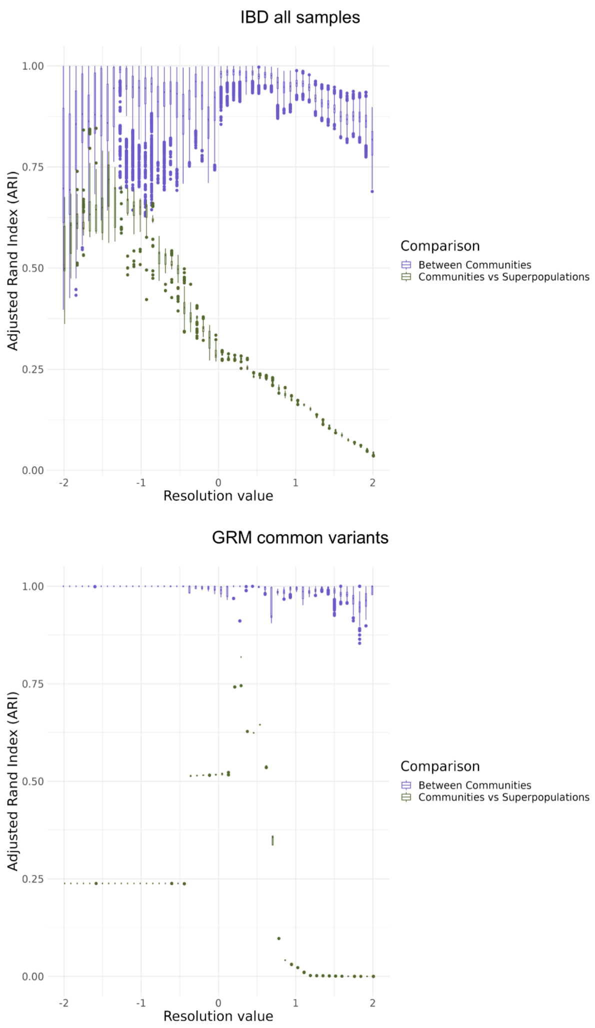


**Figure S12. Stability analysis with Adjusted Rand Index (ARI) for communities detected in the IBD without removing samples, GRM based on rare and common variants, and PCA networks at different resolutions.** The x-axis shows the resolution value. The y-axis shows the ARI value. ARI values closer to 1 indicate more individuals falling in the same communities across runs at a given resolution value. Purple boxplots summarize the comparison of community detection results across 100 independent runs at each resolution (see methods). Green boxplots represent the comparison between the independent runs and the super populations. In this case, ARI values closer to one indicate greater similarity between the detected communities and the super populations. Boxplot elements: center line, median; box limits, upper and lower quartiles; whiskers, 1.58x interquartile range; points, outliers.

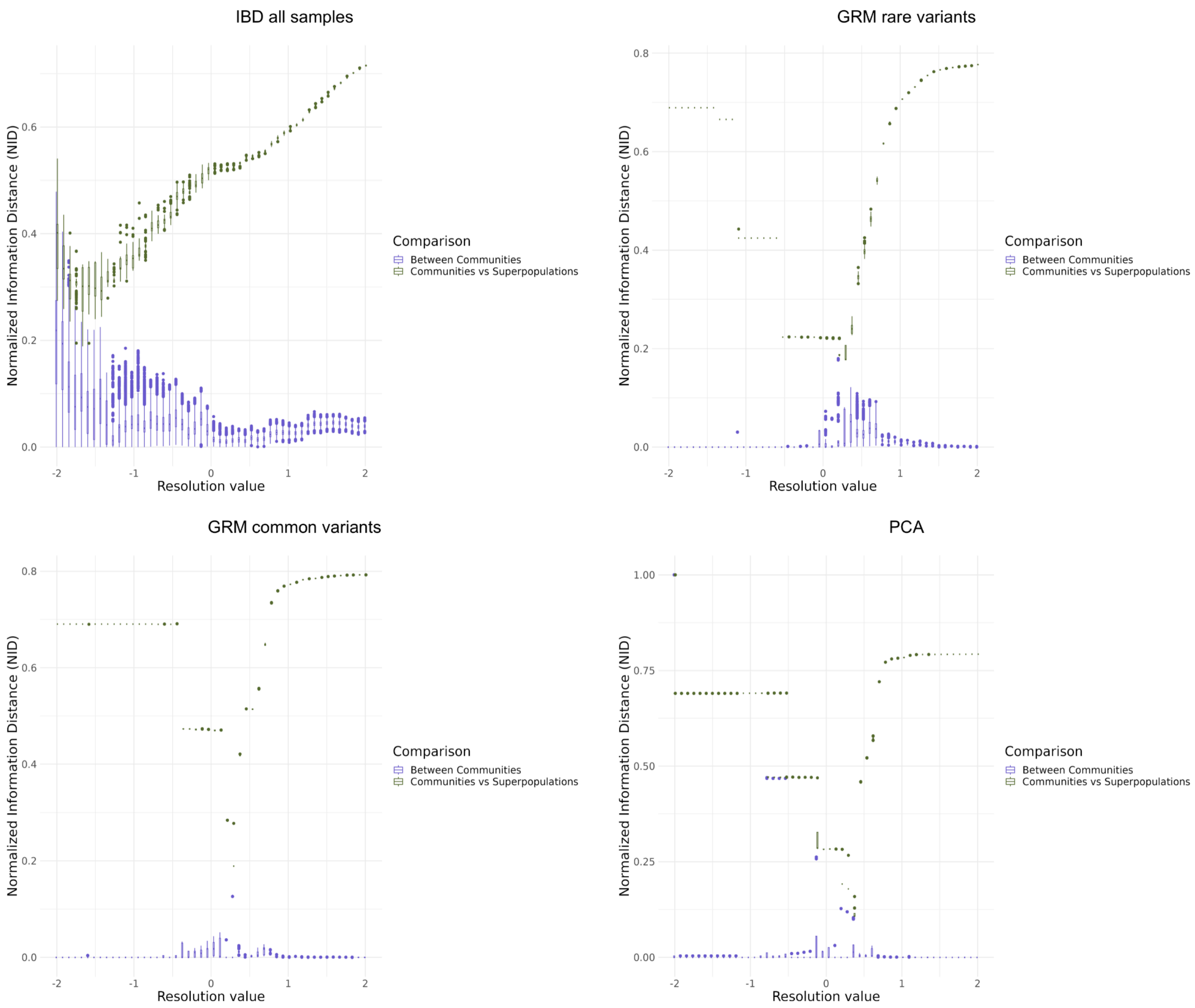


**Figure S13. Stability analysis with Normalized Information Distance (NID) for communities detected in the IBD without removing samples, GRM based on rare and common variants, and PCA networks at different resolutions.** The x-axis shows the resolution value. The y-axis shows the NID values. NID values closer to 0 indicate more individuals falling in the same communities across runs at a given resolution value. Purple boxplots summarize the comparison of community detection results across 100 independent runs at each resolution (see methods). Green boxplots represent the comparison between the independent runs and the super populations. In this case, NID values closer to 0 indicate greater similarity between the detected communities and the super populations. Boxplot elements: center line, median; box limits, upper and lower quartiles; whiskers, 1.58x interquartile range; points, outliers.

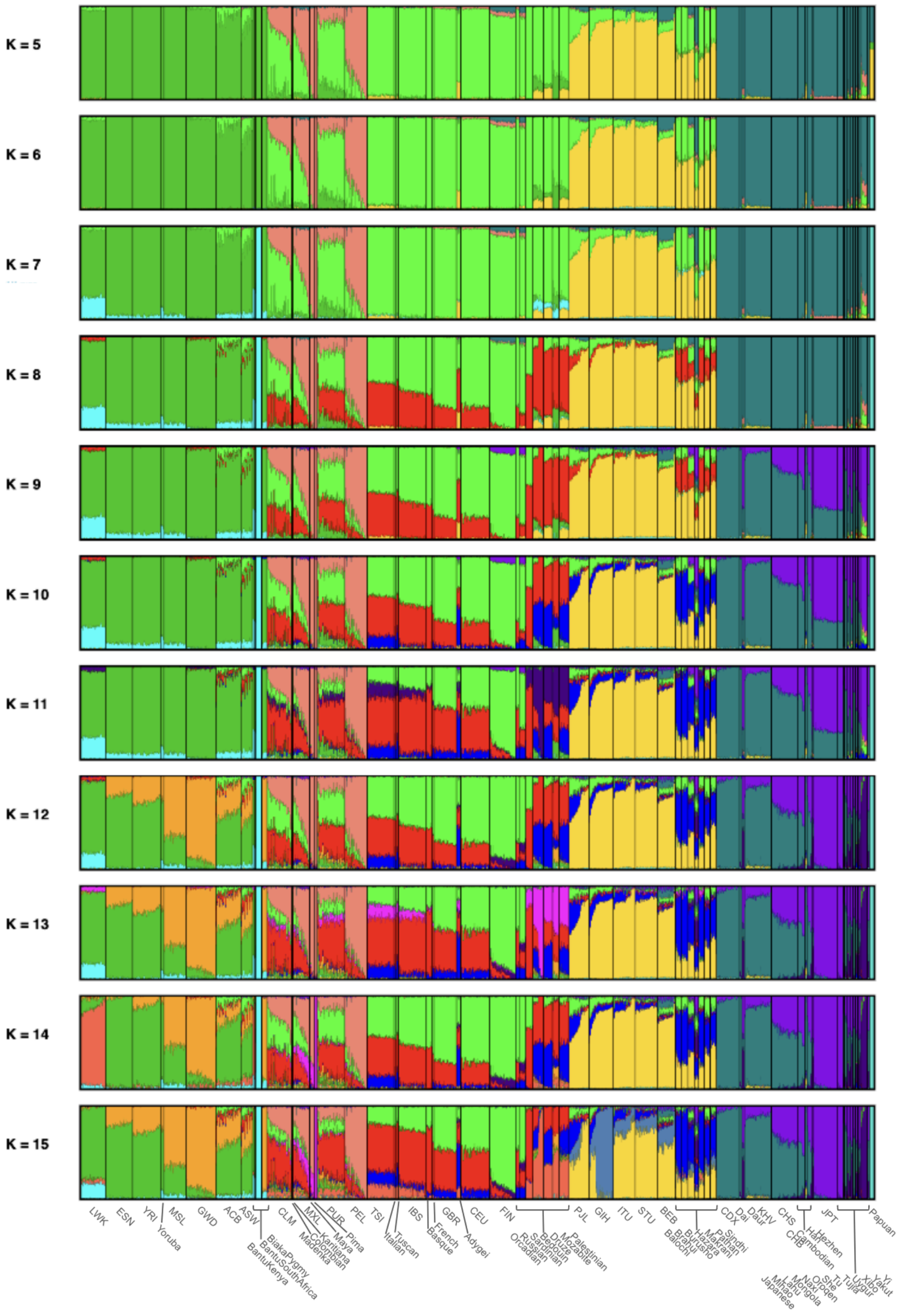


**Figure S14. Admixture analysis using the merged dataset of the 1000 Genomes Project and HGDP at K=5 to 15.** We perform admixture analysis for k values from 5 to 25 for the 2,977 individuals from the IBD network shown in the paper. This figure shows results for k = 5 to 15. Figure S16 indicates the lowest coefficient of variation (CV) error at k = 13. The analysis reveals substructural variations within continental populations and shared genetic components across different groups.

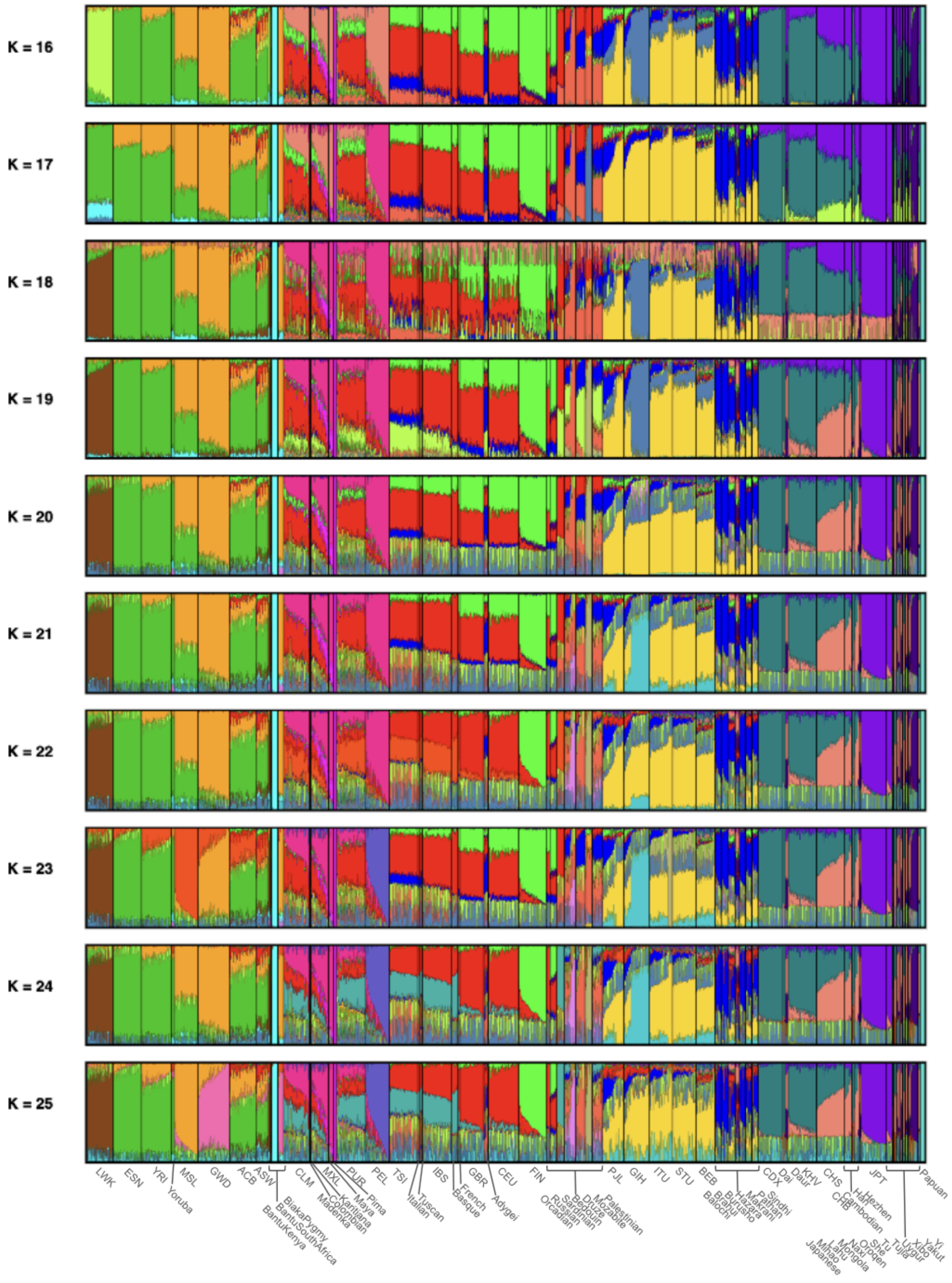


**Figure S15. Admixture analysis using the merged dataset of the 1000 Genomes Project and HGDP at K=16 to 25.** We perform admixture analysis for k values from 5 to 25 for the 2,977 individuals from the IBD network shown in the paper. This figure shows results for k = 16 to 25. Figure S16 indicates the lowest coefficient of variation (CV) error at k = 13. The analysis reveals substructural variations within continental populations and shared genetic components across different groups.

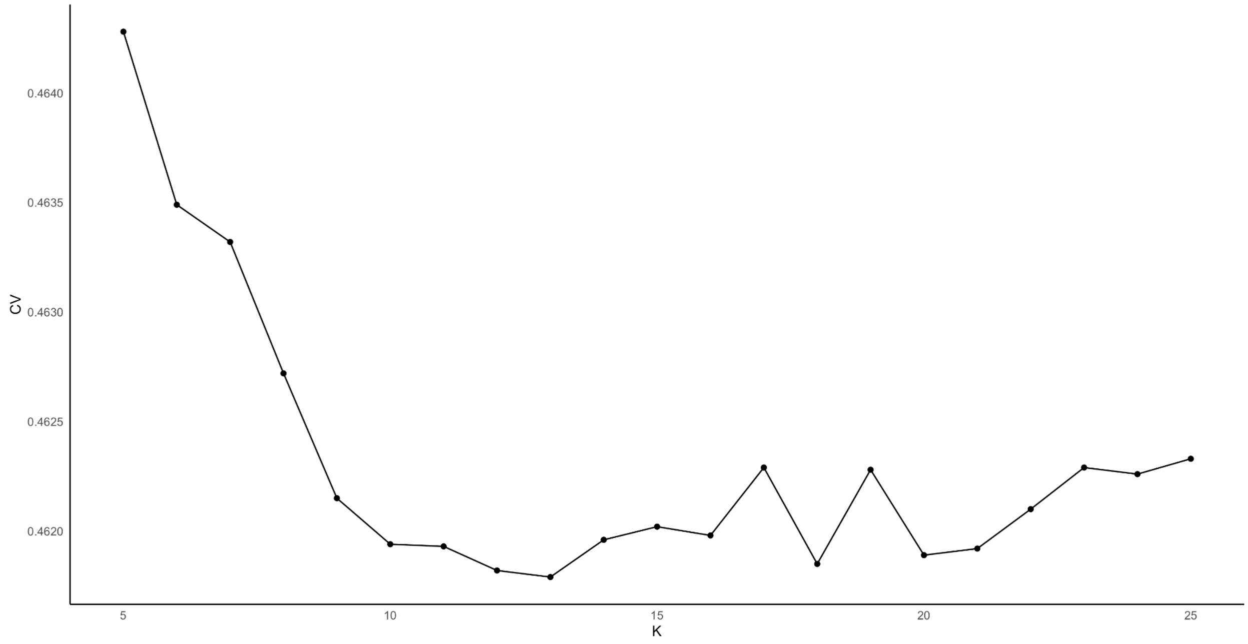


**Figure S16. Coefficient of variation (CV) Error for different values of K in the ADMIXTURE analysis.** This figure illustrates the coefficient of variation (CV) resulting from the admixture analysis of 2,977 individuals from the IBD network shown in the study, across a range of k values from 5 to 25. The X-axis denotes the k values, while the Y-axis displays the associated CV errors. The minimum CV error was observed at k=13.

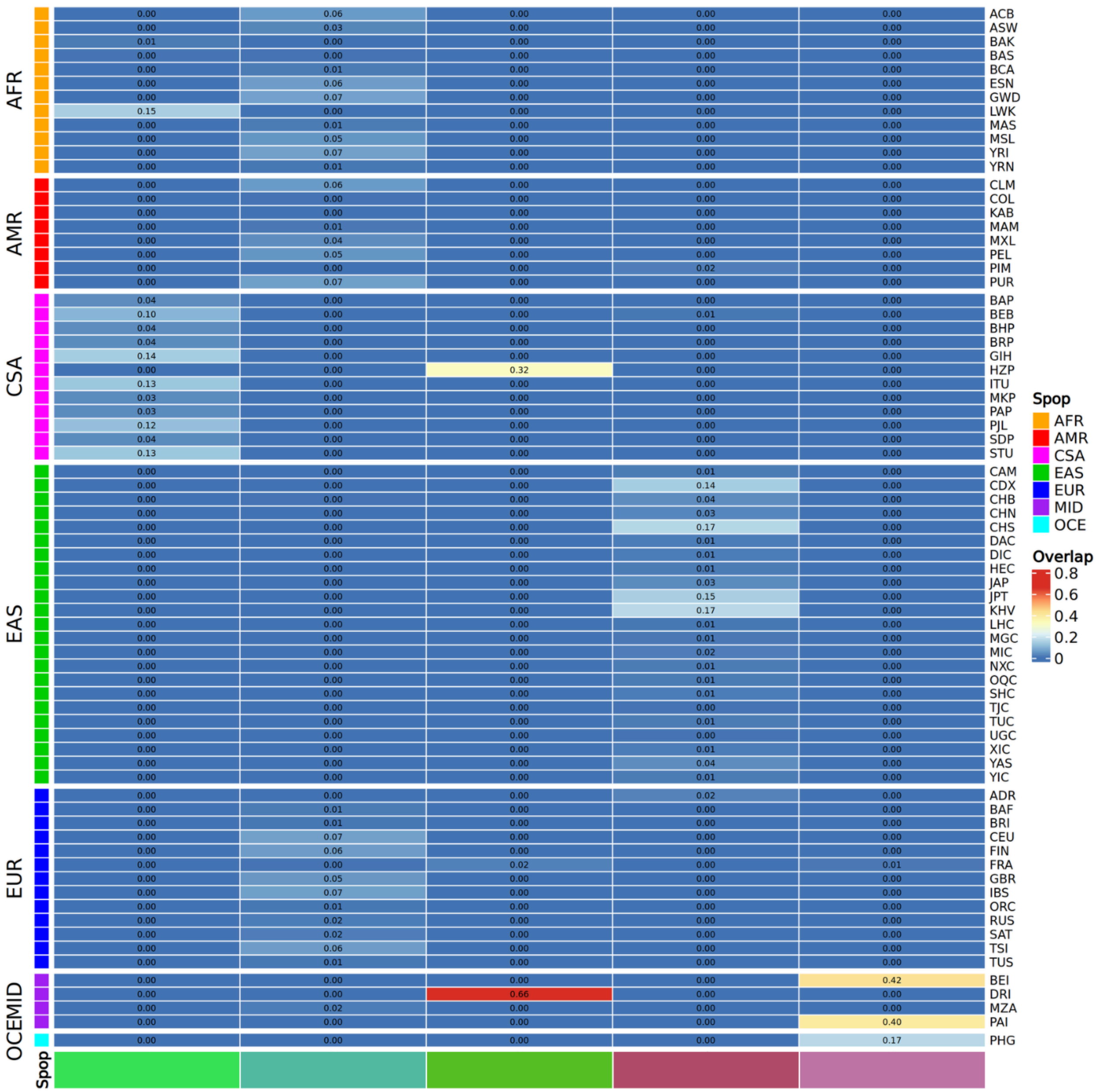


**Figure S17. Overlap heatmap between communities at resolution value -2 and cohorts from 1000G and HGDP**. Each row represents a population grouped by super populations according to the tab on the left, and each column represents a community of a given color in the tab below. The cells contain the proportion of individuals from each population that make up each community. Therefore, if we add them up, each column should equal 1.

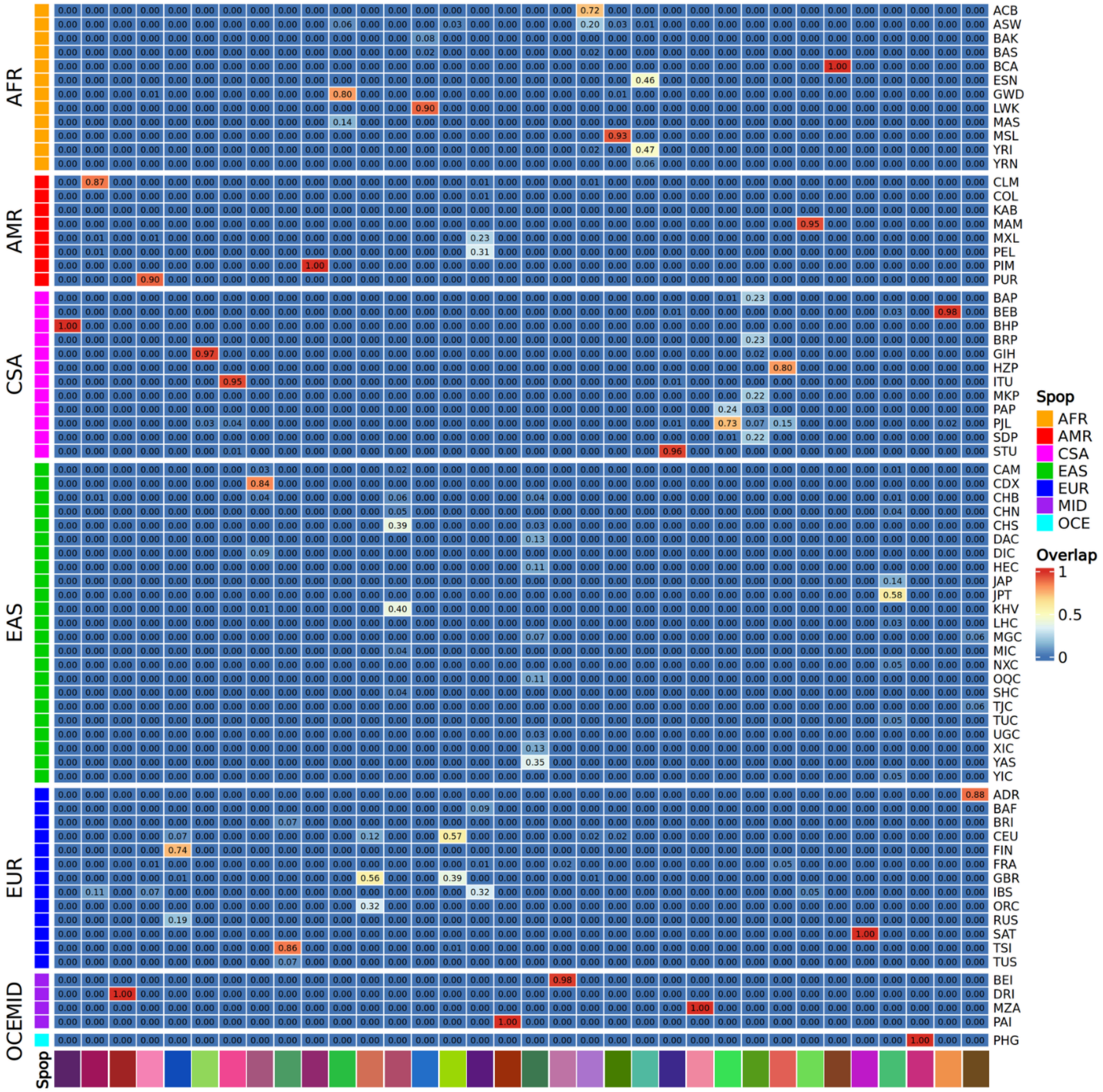


**Figure S18. Overlap heatmap between communities at resolution value -0.041 and cohorts from 1000G and HGDP**. Each row represents a population grouped by super populations according to the tab on the left, and each column represents a community of a given color in the tab below. The cells contain the proportion of individuals from each population that make up each community. Therefore, if we add them up, each column should equal 1.

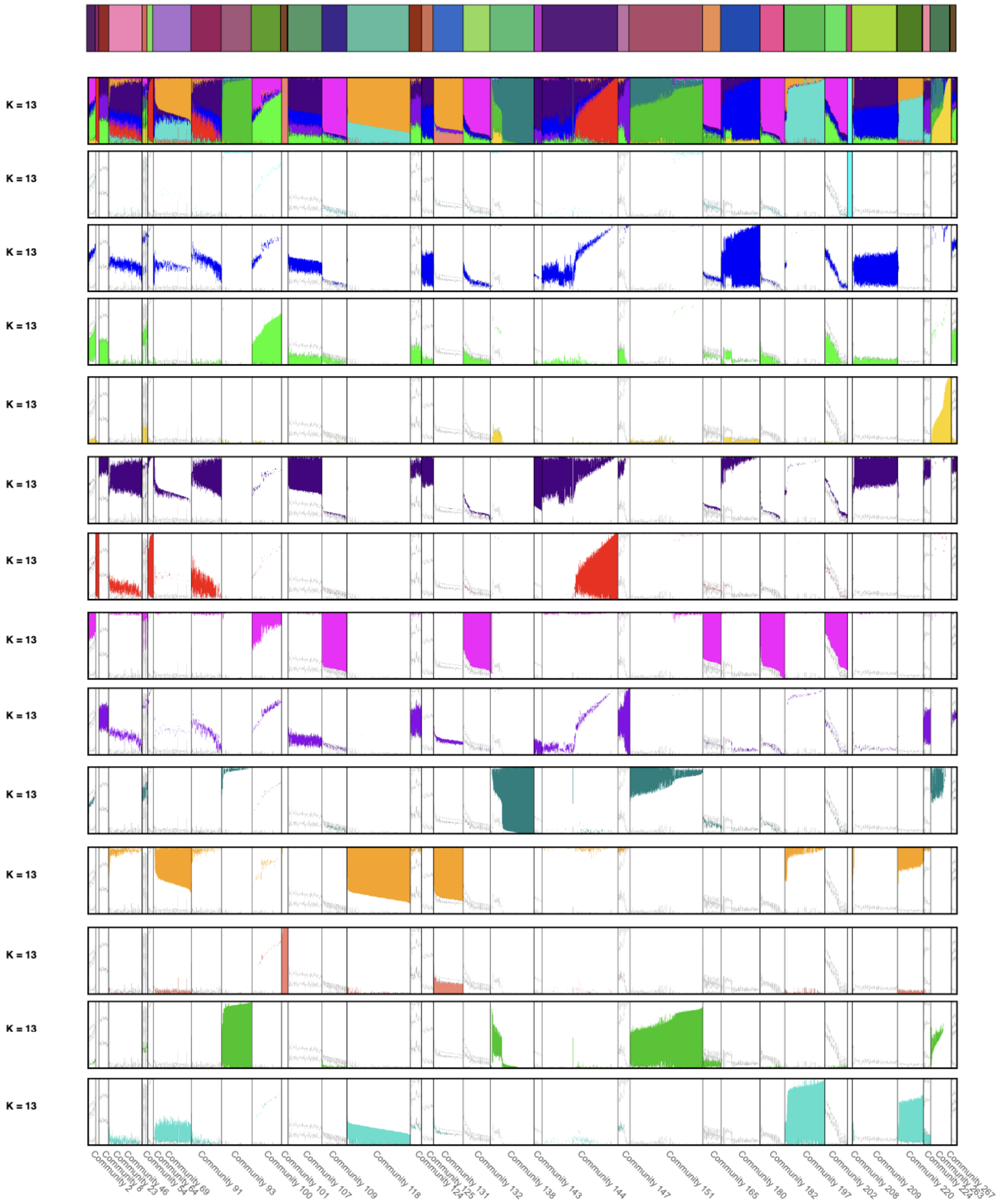


**Figure S19. Components detected in the ADMIXTURE analysis at K=13 across the communities.** A resolution of -0.041 is considered. Each plot highlights the detected components individually. None of the components are found exclusively in a single community.

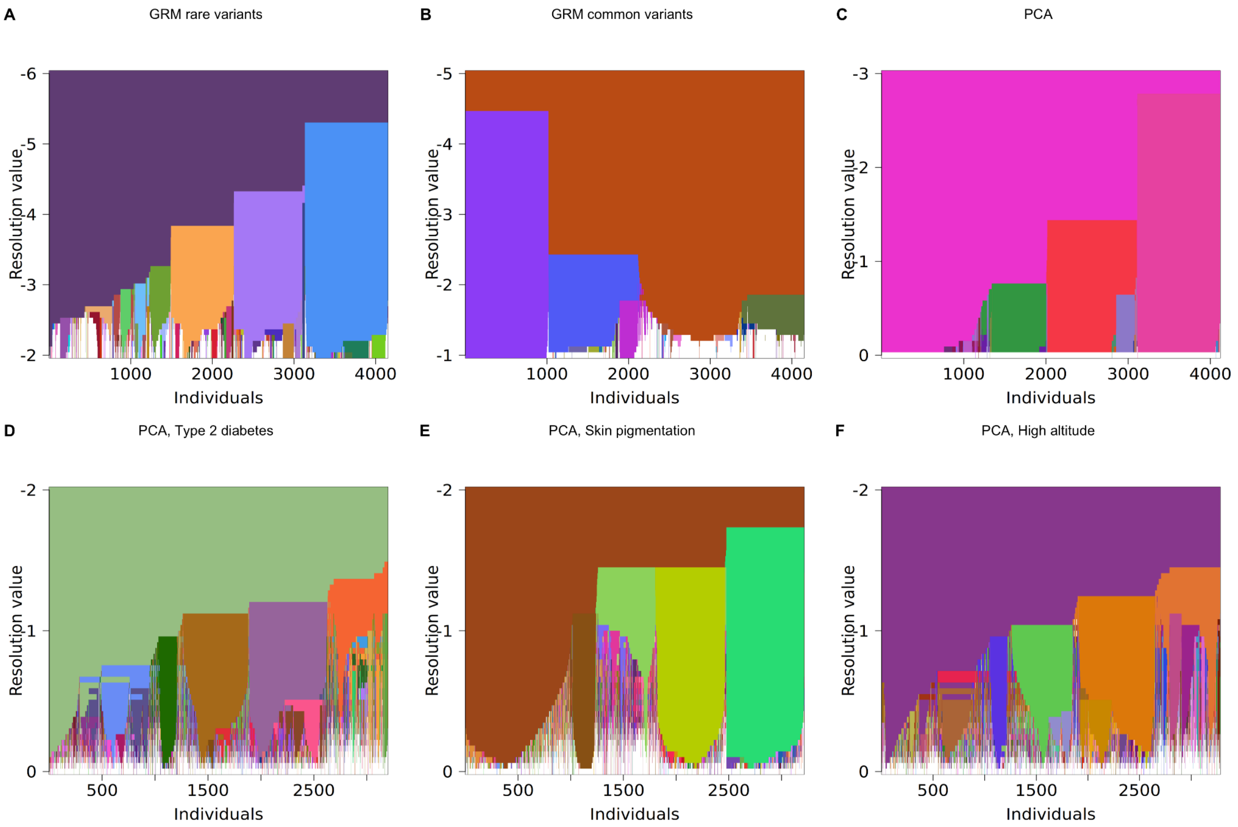


**Figure S20.** ***Resolution plots*** **(Leiden algorithm) from networks using different definitions of genetic similarity and different subsets of genetic variants reveal different aspects of genetic relatedness.** *Resolution plots* summarize community detection results at 50 resolution values; Communities that do not have more than 6 members in any resolution are colored in white. The x-axis represents the individuals and the y-axis corresponds to the resolution value A) *Resolution plot* for the network based on the Genetic Relationship Matrix (GRM) estimated on rare variants (n=4,150). B) *Resolution plot* for the network based on the GRM estimated on common variants (n=4,150). C) *Resolution plot* for the network based on Principal Component Analysis (PCA) correlation (n=4,119). *Resolution plots* for trait-PCA-based networks using only independent variants in: D) Type 2 diabetes associated genes (n= 3,199; 15 genes)[36]. E) Skin pigmentation associated genes (n=3,214; 38 genes)[37]. F) Genes associated with or inferred to be under natural selection for Altitude adaptation (n=3,281; 7 genes)[38].

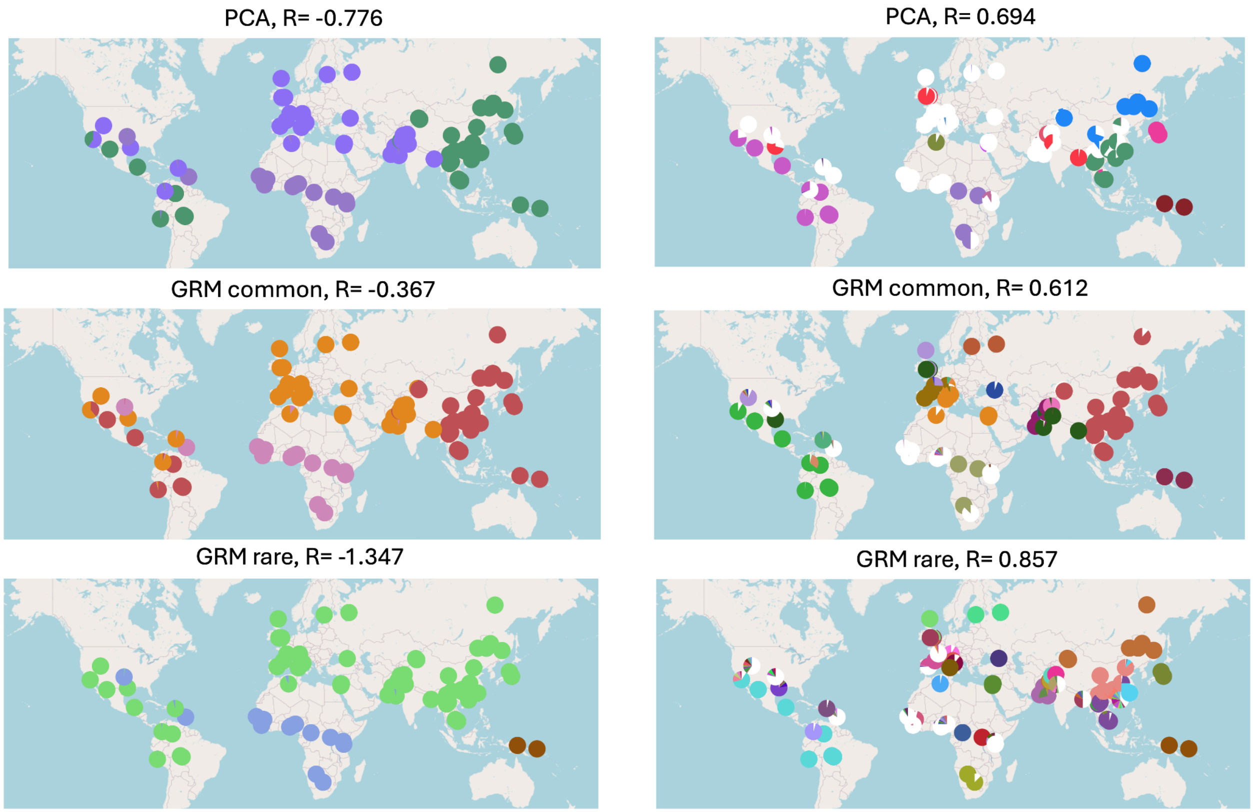


**Figure S21. Geographic distribution of communities based on GRM or PCA.** Maps on the left side show the first three communities that emerge from the data. The panels on the right show the differences in community sizes detected by the different metrics. R values were chosen to display communities before they fragment into very small groups (<6 members).

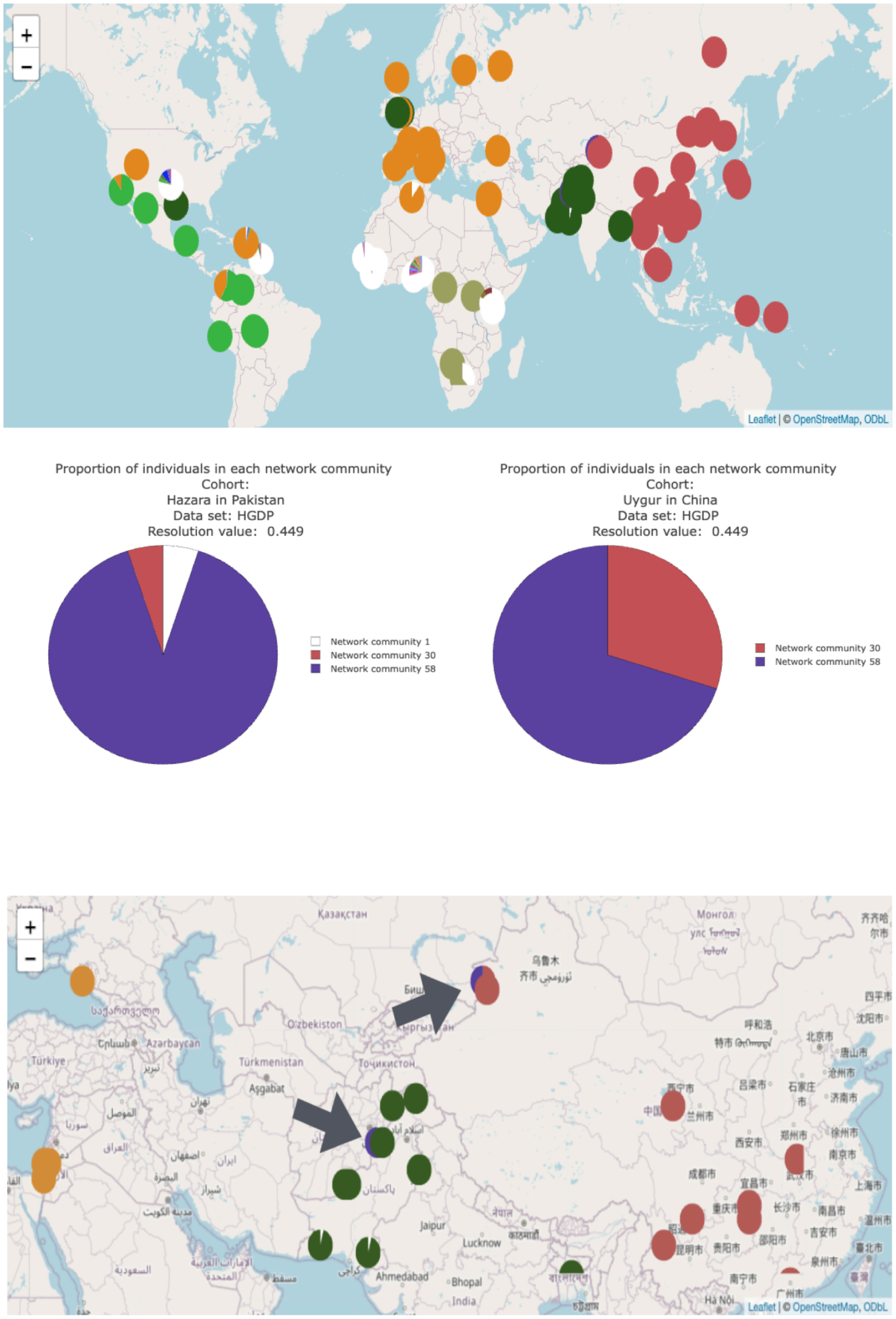


**Figure S22. Community formation in Hazara and Uygur Cohorts with increasing resolution in the GRM (common variants).** The geographic distribution and proportion of individuals in each community for the Hazara cohort in Pakistan and the Uygur cohort in China, as determined by the Genetic Relationship Matrix (GRM) based on common variants at a resolution of 0.449. In the below section of the figure, a zoomed-in section of the map is displayed, with arrows indicating the locations of the aforementioned cohorts. A high proportion of Hazara and Uygur individuals group into the same community according to the GRM (common variants).

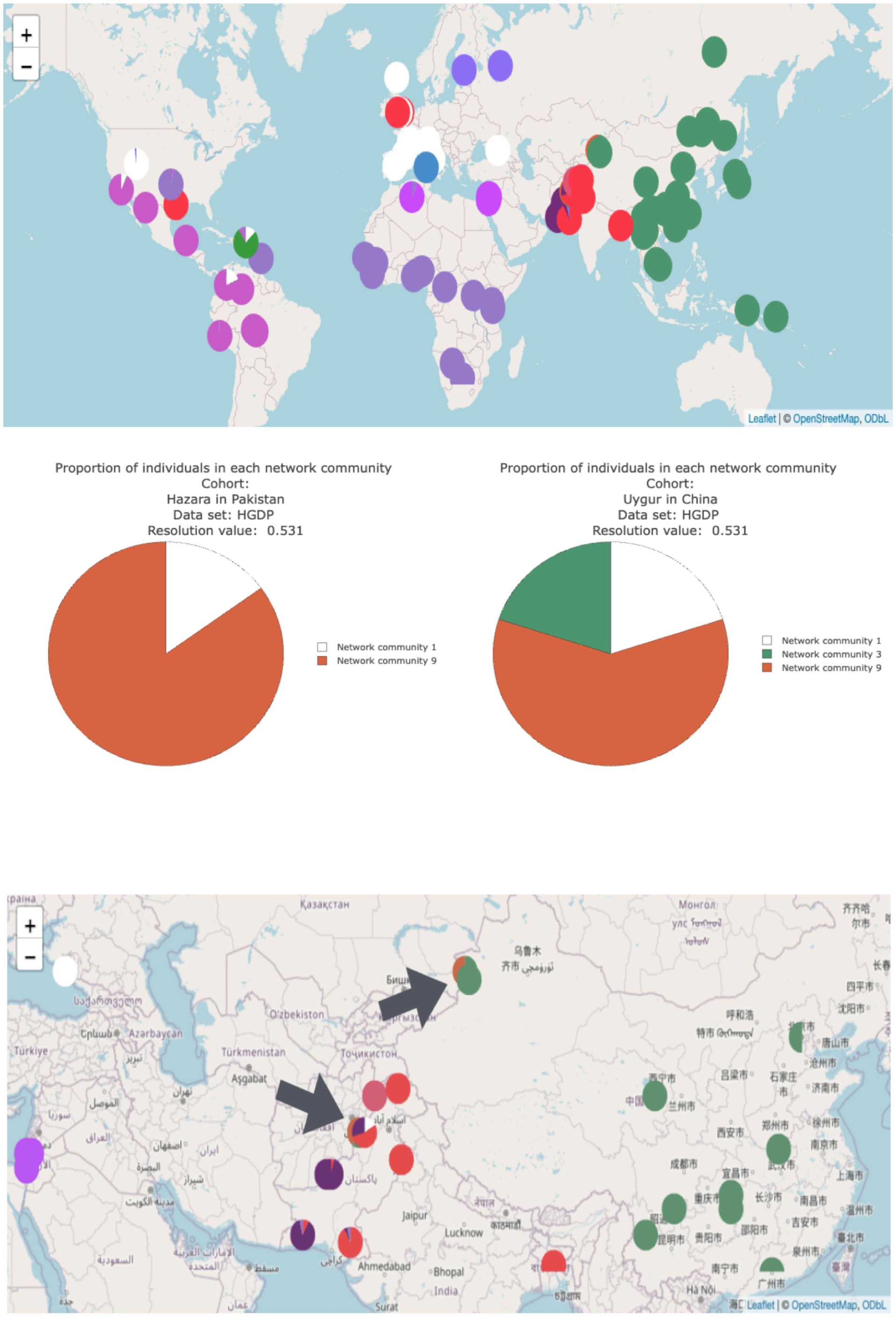


**Figure S23. Community formation in Hazara and Uygur Cohorts with increasing resolution in the PCA.** The geographic distribution and proportion of individuals in each community for the Hazara cohort in Pakistan and the Uygur cohort in China according to the Principal Components Analysis (PCA) at a resolution of 0.531. In the below section of the figure, a zoomed-in section of the map is displayed, with arrows indicating the locations of the aforementioned cohorts. A high proportion of Hazara and Uygur individuals group into the same community according to the PCA.

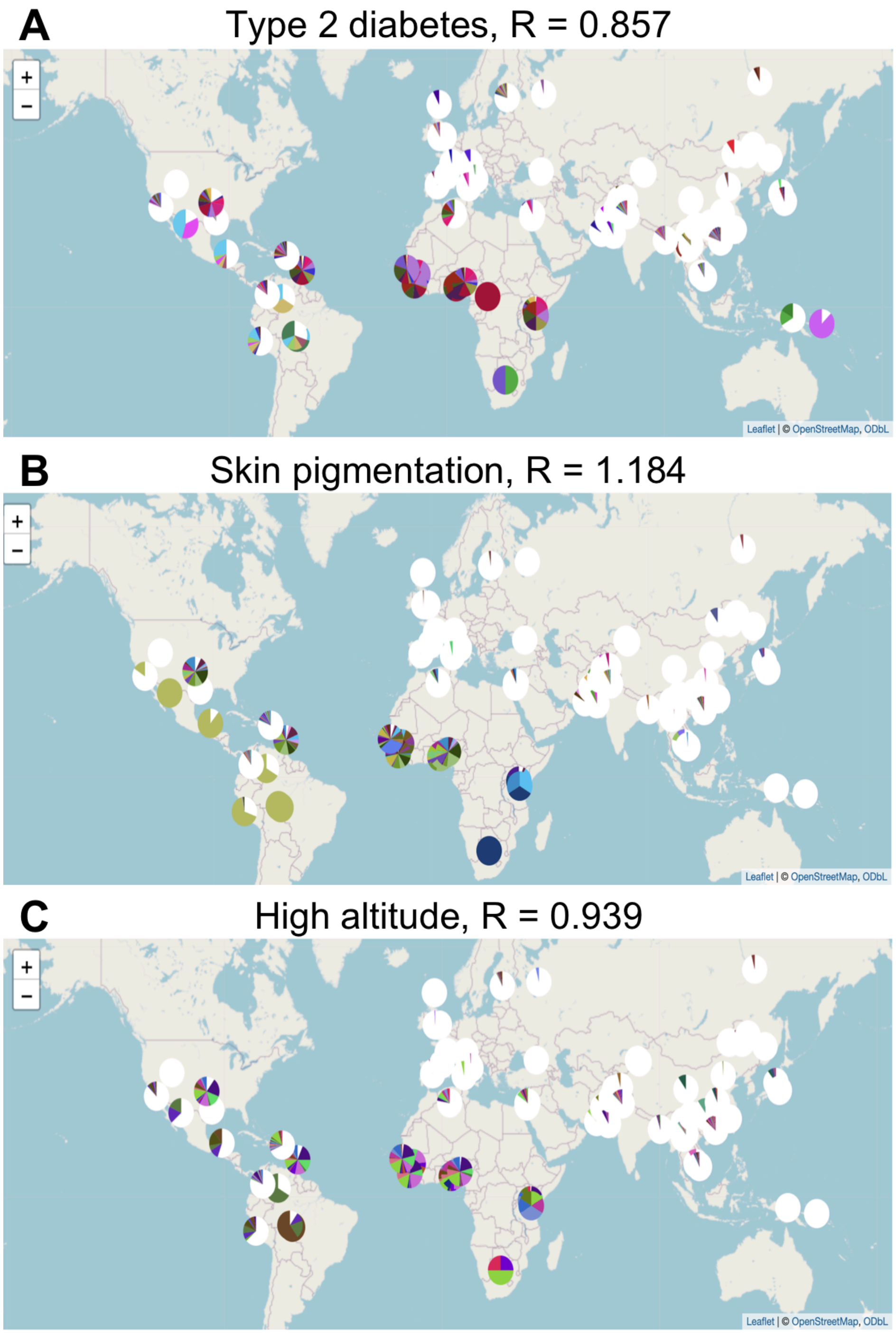


**Figure S24. Substructure in Africa detected on trait-specific results for the PCA networks. A)** Using variants involved in Type 2 diabetes at a resolution of 0.857. **B)** Using skin pigmentation associated genes at a resolution of 1.184. **C)** With respect to genes associated with or inferred to be under natural selection for Altitude adaptation at a resolution of 0.939. In all situations, at higher resolutions, a greater substructure of communities was detected, as observed in Africa.

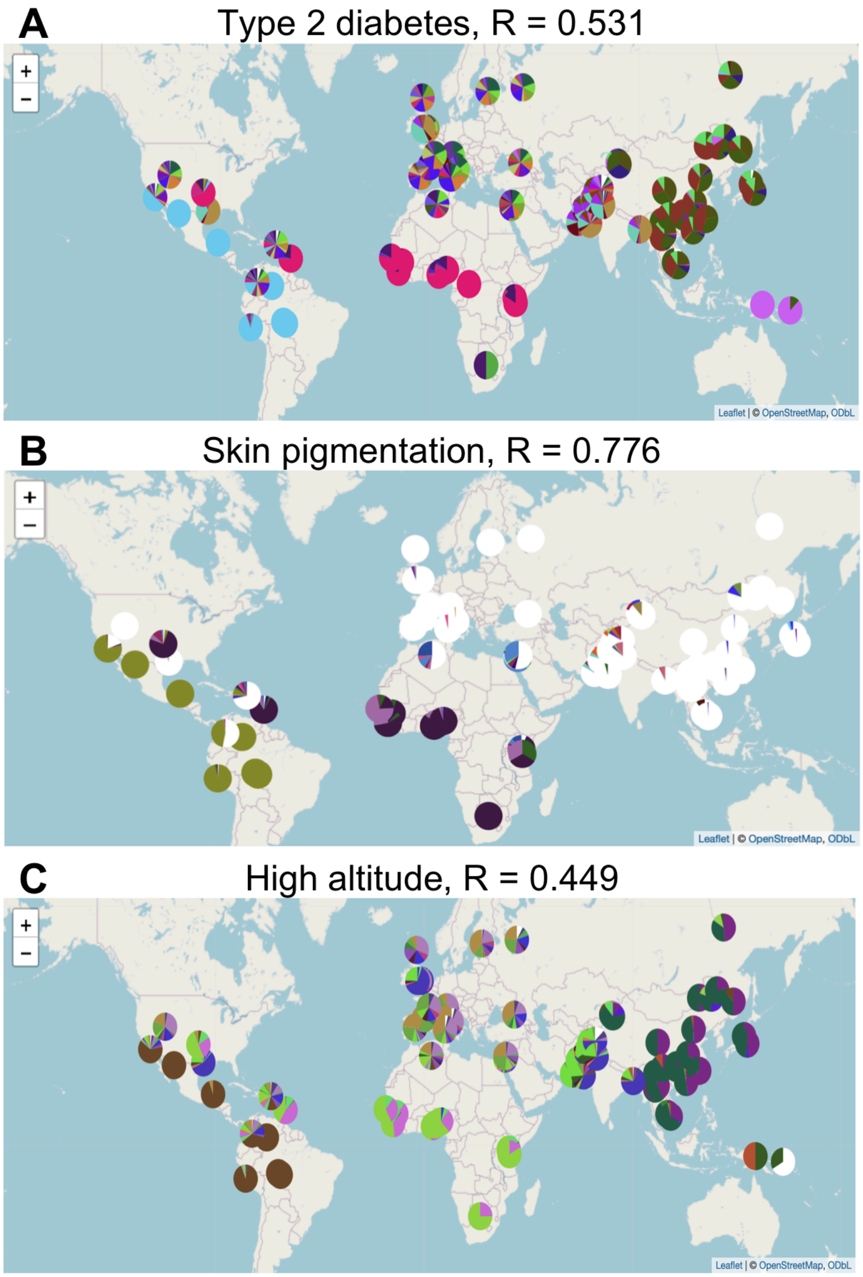


**Figure S25. The Americas in the trait-specific results for the PCA networks**. **A)** Using variants involved in Type 2 diabetes at a resolution of 0.531. **B)** Using skin pigmentation associated genes at a resolution of 0.776. **C)** Genes associated with or inferred to be under natural selection for Altitude adaptation at a resolution of 0.449. As the resolution increases, indigenous individuals from the Americas, as well as some individuals in urban contexts (Colombians, Peruvians, Mexicans in Los Angeles), consistently remain in the same community.

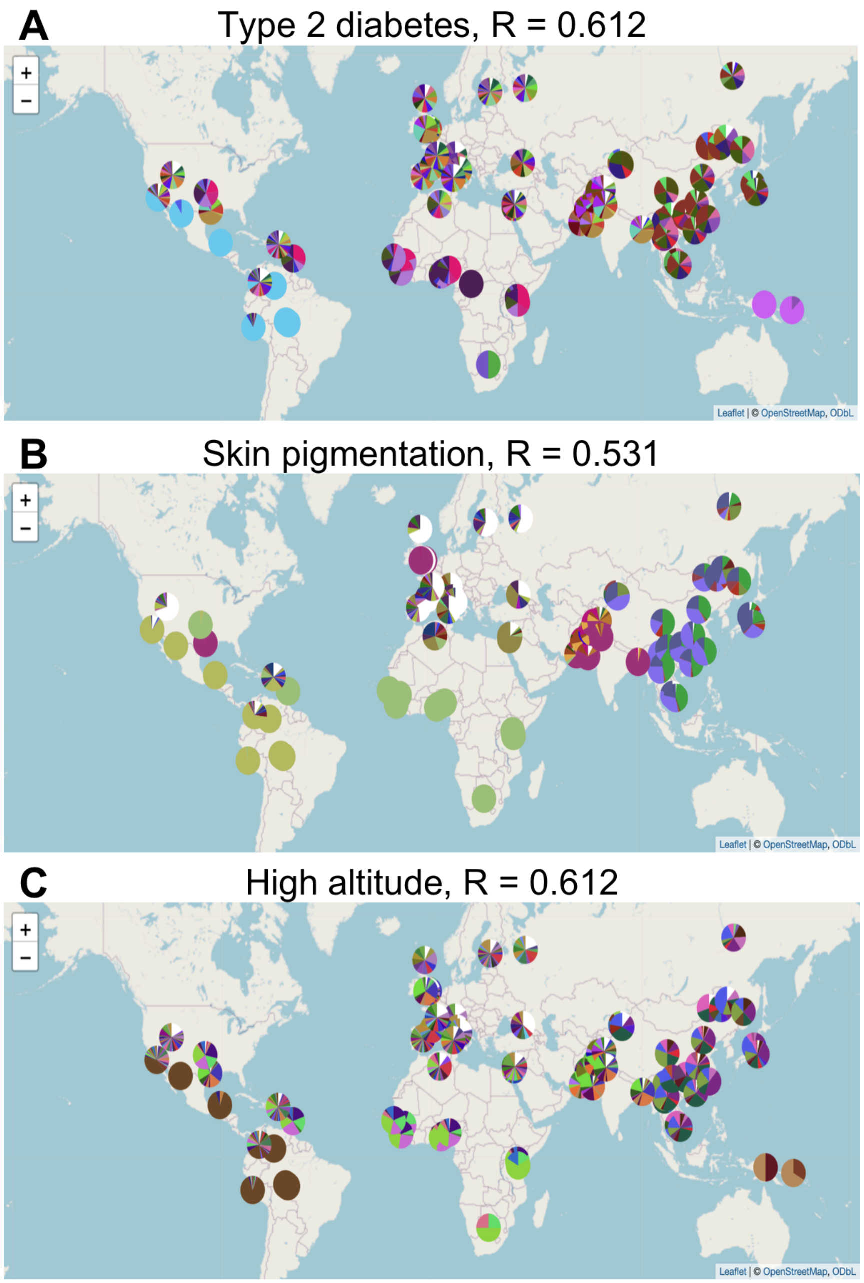


**Figure S26. Substructure Variation Across Genomic Regions Associated with Different Traits. A)** Using variants involved in Type 2 diabetes at a resolution of 0.612. **B)** Using skin pigmentation associated genes at a resolution of 0.531. **C)** Genes associated with or inferred to be under natural selection for Altitude adaptation at a resolution of 0.612. Substructure observed is not limited to defined cohorts.

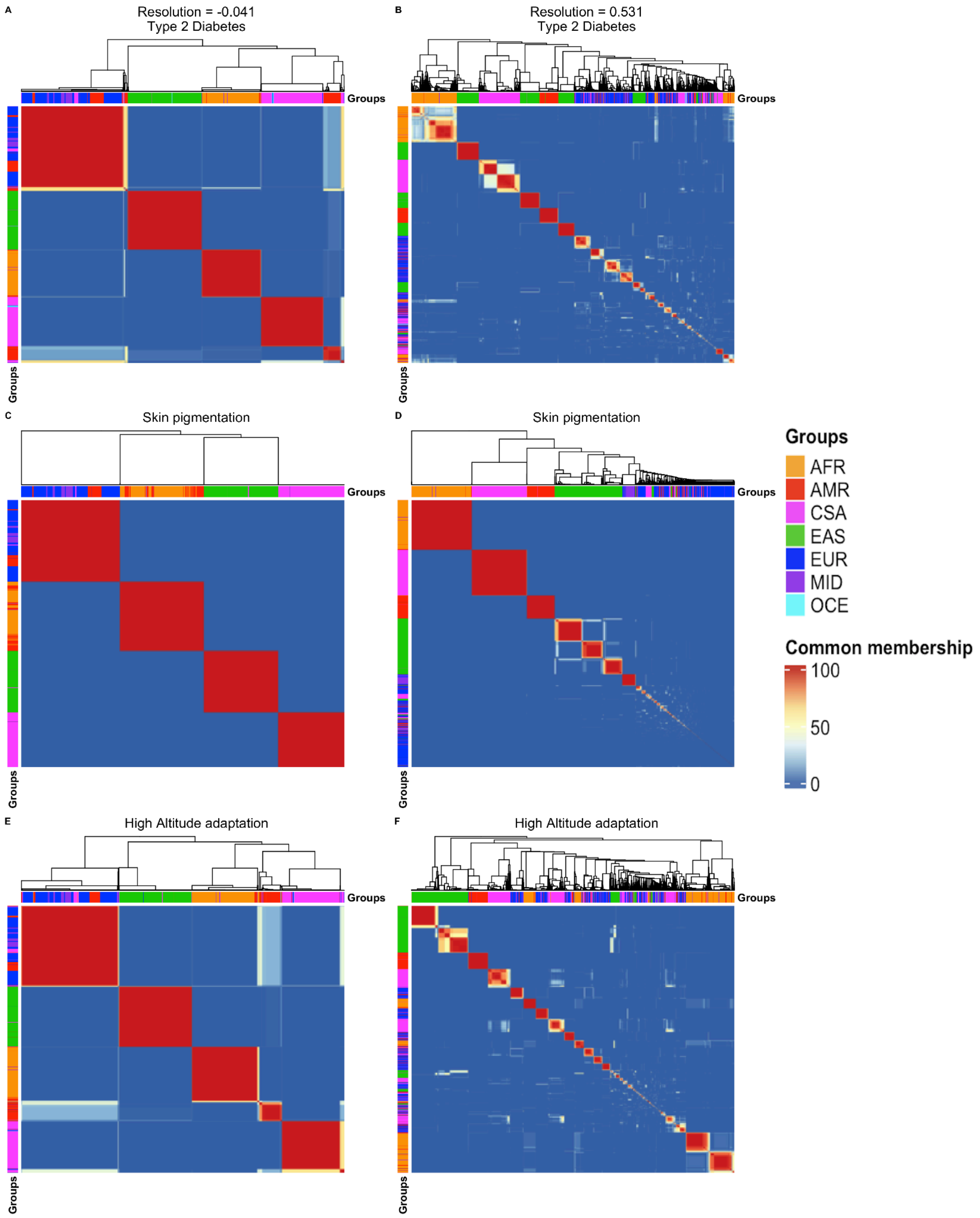


**Figure S27. Heatmaps of shared membership in trait-specific PCA communities. A,B)** Using variants involved in Type 2 diabetes at a resolution of -0.041 and 0.531, respectively. **C,D)** Using skin pigmentation associated genes at a resolution of -0.041 and 0.531, respectively. **E,F)** According to genes associated with or inferred to be under natural selection for Altitude adaptation at a resolution of -0.041 and 0.531, respectively. Along the X and Y axes, individuals that consistently cluster together are hierarchically grouped and they are indicated according to the continental group to which they belong. A warm color indicates a higher level of shared membership. The distribution of shared membership exhibits greater similarity at a resolution of -0.041 across the three specific traits studied.

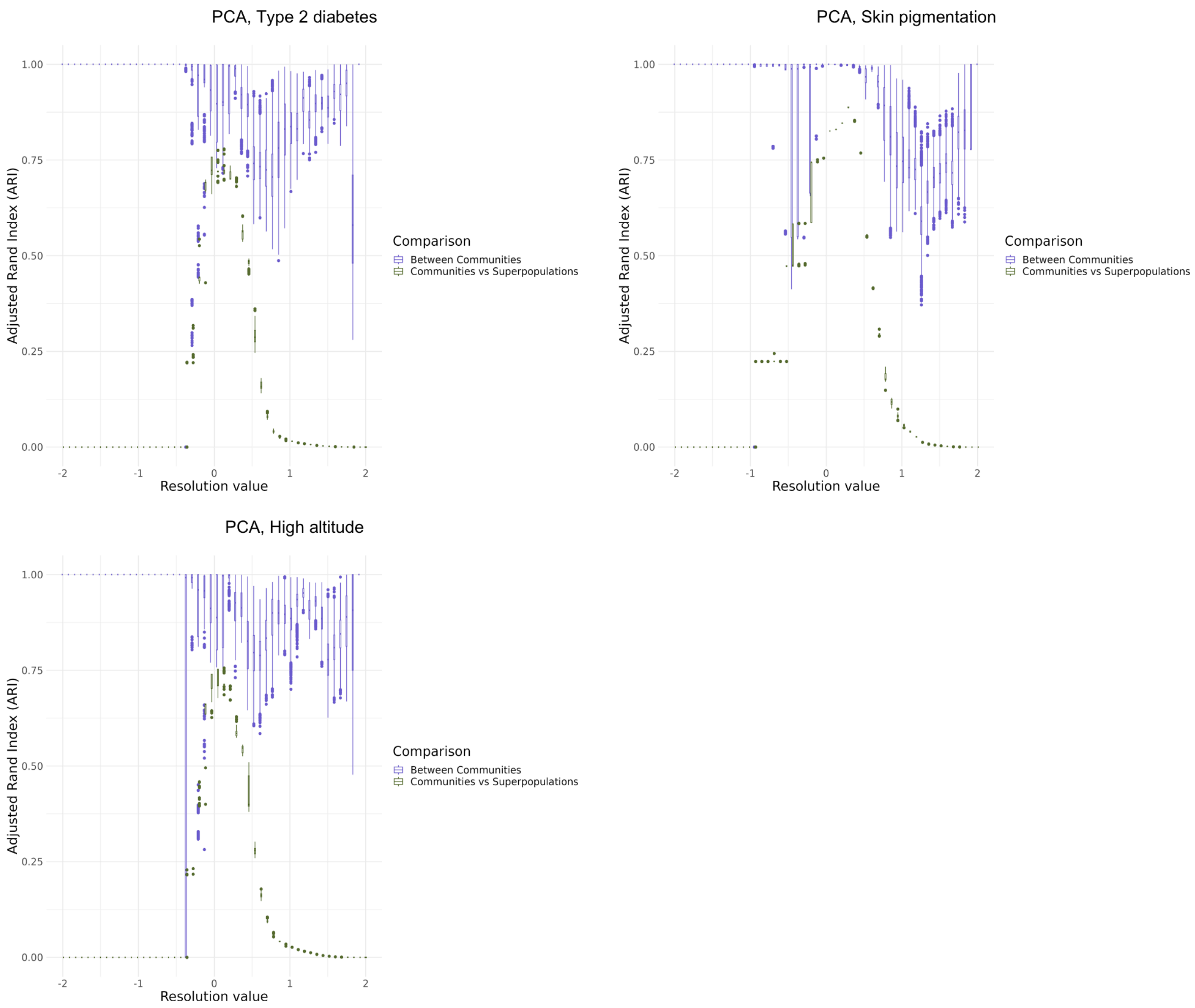


**Figure S28. Stability of communities for PCA results for Type 2 diabetes, Skin pigmentation, and High altitude at any given resolution is shown with Adjusted Rand Index (ARI).** The x-axis shows the resolution value. The y-axis shows the ARI value. Each boxplot summarizes the results for 100 runs (see methods). ARI values closer to 1 indicate more individuals falling in the same communities across runs at a given resolution value. Purple boxplots summarize the comparison of community detection results across 100 independent runs at each resolution (see methods). Green boxplots represent the comparison between the independent runs and the super populations. In this case, ARI values closer to one indicate greater similarity between the detected communities and the super populations. Boxplot elements: center line, median; box limits, upper and lower quartiles; whiskers, 1.58x interquartile range; points, outliers.

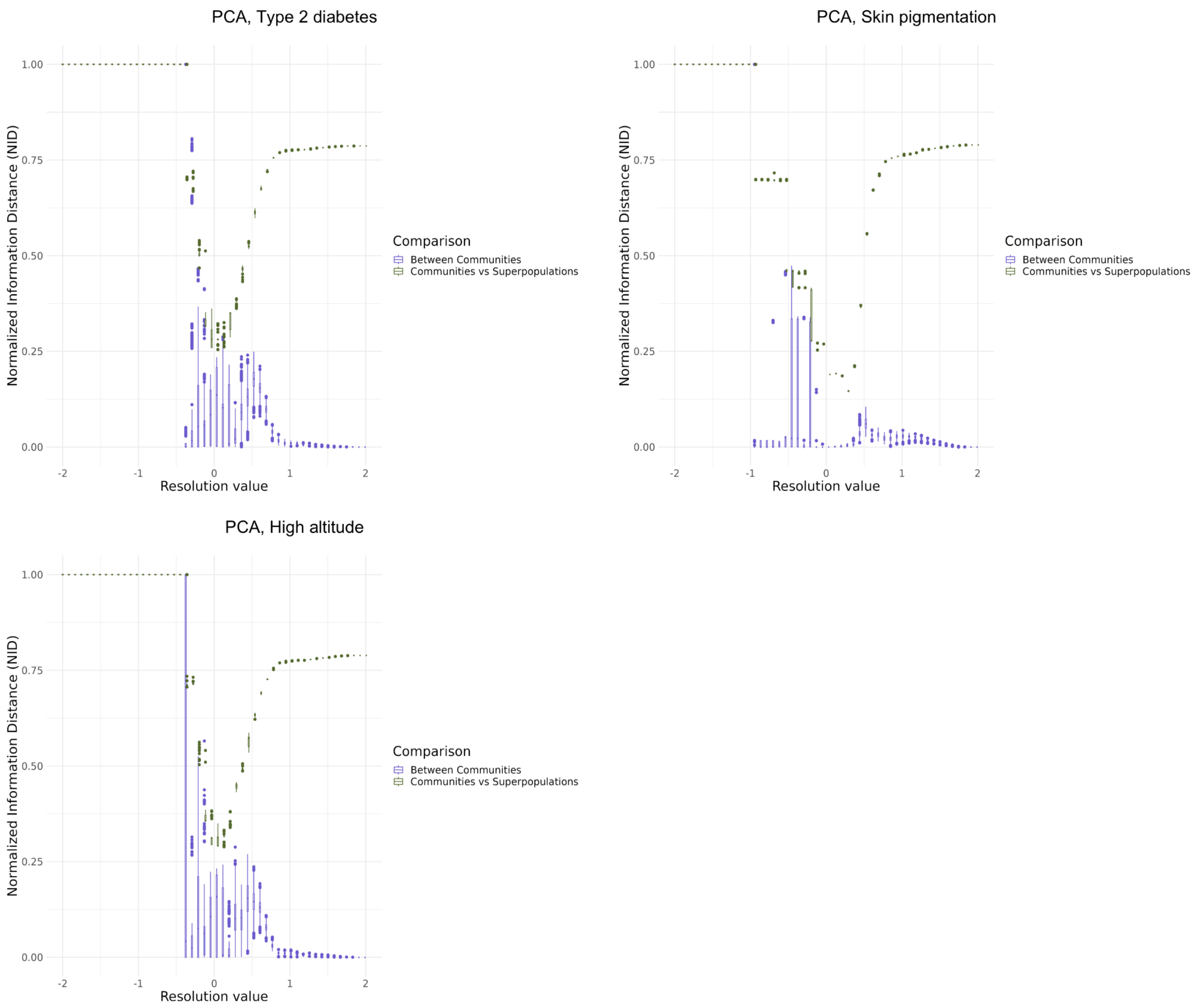


**Figure S29. Stability of communities for PCA results for Type 2 diabetes, Skin pigmentation, and High altitude at any given resolution is shown with Normalized Information Distance (NID).** The x-axis shows the resolution value. The y-axis shows the NID values. NID values closer to 0 indicate more individuals falling in the same communities across runs at a given resolution value. Purple boxplots summarize the comparison of community detection results across 100 independent runs at each resolution (see methods). Green boxplots represent the comparison between the independent runs and the super populations. In this case, NID values closer to 0 indicate greater similarity between the detected communities and the super populations. Boxplot elements: center line, median; box limits, upper and lower quartiles; whiskers, 1.58x interquartile range; points, outliers.

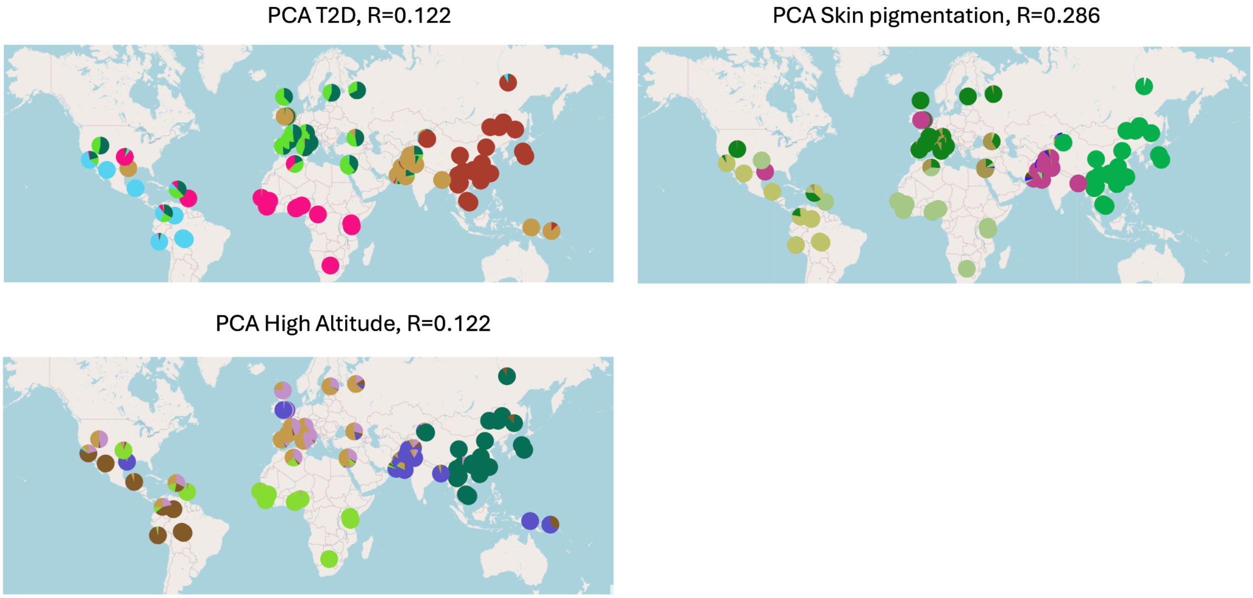


**Figure S30. Geographic distribution of communities at the resolution with the highest median coincidence with super populations (PCA trait specific).** Geographic distribution of detected communities for PCA trait specific (Type 2 diabetes, High Altitude adaptations, and skin pigmentation) at the resolution where the highest median ARI is observed when comparing communities with super populations.

### Supplementary Table Captions

**Supplementary Table 1. IBD segments shared pairwise.** Sum of the length of the IBD segments (>5cM) shared pairwise between the individuals in the jointly called dataset 1000G + HGDP.

**Supplementary Table 2. Sample metadata and inclusion on the different metrics.** Metadata for each individual including cohort, super population, as well as sampling location of the cohort. Other columns indicate if the individual was included in the different metric analysis.

**Supplementary Table 3. Cohort sizes for the different metrics.** Cohort sizes for each of the different metrics analyzed.

**Supplementary Table 4. Wilcoxon results for community detection based on IBD.** Wilcoxon test results for IBD (Bonferroni threshold = 0.001). The data includes median and variance values for comparisons between communities, and between communities and super populations; also shown is the range of detected community counts at each resolution.

**Supplementary Table 5. Wilcoxon results for community detection based on PCA.** Wilcoxon test results for PCA (Bonferroni threshold = 0.001). The data includes median and variance values for comparisons between communities, and between communities and super populations; also shown is the range of detected community counts at each resolution.

**Supplementary Table 6. Wilcoxon results for community detection based on GRM common variants.** Wilcoxon test results for GRM common variants (Bonferroni threshold = 0.001). The data includes median and variance values for comparisons between communities, and between communities and super populations; also shown is the range of detected community counts at each resolution.

**Supplementary Table 7. Wilcoxon results for community detection based on GRM rare variants.** Wilcoxon test results for GRM rare variants (Bonferroni threshold = 0.001). The data includes median and variance values for comparisons between communities, and between communities and super populations; also shown is the range of detected community counts at each resolution.

**Supplementary Table 8. Wilcoxon results for community detection based on trait-specific analysis (PCA skin pigmentation).** Wilcoxon test results for PCA at each resolution, focusing on genes associated with skin pigmentation (Bonferroni threshold = 0.001). The data includes median and variance values for comparisons between communities, and between communities and super populations; also shown is the range of detected community counts at each resolution.

**Supplementary Table 9. Wilcoxon results for community detection based on trait-specific analysis (PCA high altitude adaptation).** Wilcoxon test results for PCA at each resolution, focusing on genes associated with adaptation to high altitude (Bonferroni threshold = 0.001). The data includes median and variance values for comparisons between communities, and between communities and super populations; also shown is the range of detected community counts at each resolution.

**Supplementary Table 10. Wilcoxon results for community detection based on trait-specific analysis (PCA Type 2 diabetes).** Wilcoxon test results for PCA at each resolution, focusing on genes associated with T2D (Bonferroni threshold = 0.001). The data includes median and variance values for comparisons between communities, and between communities and super populations; also shown is the range of detected community counts at each resolution.
