## Extended Data Table 1 for "The multi-scale complexity of human genetic variation beyond continental groups"

| High Altitude Adaptation | | | |
|---|---|---|---|
| **Gene** | **Chromosome** | **Start (bp)** | **End(bp)** |
| EPAS1 | 2 | 46293667 | 46386697 |
| EGLN1 | 1 | 231363751 | 231422287 |
| PPARA | 22 | 46150521 | 46243755 |
| CBARA1 | 10 | 72367340 | 72626131 |
| VAV3 | 1 | 107571161 | 107965180 |
| ARNT2 | 15 | 80404350 | 80597933 |
| THRB | 3 | 24117153 | 24495756 |
