## Extended Data Table 2 for "The multi-scale complexity of human genetic variation beyond continental groups"

| Skin pigmentation | | | | | | | |
|---|---|---|---|---|---|---|---|
| **Gene** | **Chr** | **Start (bp)** | **End(bp)** | **Gene** | **Chr** | **Start (bp)** | **End(bp)** |
| OCA2 | 15 | 27754875 | 28099315 | WNT1 | 12 | 48978322 | 48982620 |
| SLC24A5 | 15 | 48120990 | 48142672 | SILV | 12 | 55954105 | 55973317 |
| SLC45A2 | 5 | 33944623 | 33984693 | OPRM1 | 6 | 154010946 | 154246867 |
| TYR | 11 | 89177875 | 89295759 | EGFR | 7 | 55019017 | 55211628 |
| MFSD12 | 19 | 3538261 | 3574290 | ZNF804B | 7 | 88759700 | 89338528 |
| DDB1 | 11 | 61299451 | 61342596 | PDE4B | 1 | 65792514 | 66374579 |
| TMEM138 | 11 | 61361964 | 61377890 | RIPK5 | 1 | 205142505 | 205211702 |
| HERC2 | 15 | 28111040 | 28322179 | PA2G4P4 | 3 | 156809551 | 156810732 |
| IRF4 | 6 | 391739 | 411443 | PPARGC1B | 5 | 149730298 | 149855022 |
| BEND7 | 10 | 13438484 | 13529014 | AHR | 7 | 16916359 | 17346152 |
| PRPF18 | 10 | 13586939 | 13668445 | AGR3 | 7 | 16859412 | 16881987 |
| MC1R | 16 | 89912119 | 89920973 | TRPS1 | 8 | 115408496 | 115809673 |
| ASIP | 20 | 34194569 | 34269344 | BNC2 | 9 | 16409503 | 16870843 |
| TYRP1 | 9 | 12685439 | 12710285 | EMX2 | 10 | 117542445 | 117549546 |
| SMARCA2 | 9 | 1980290 | 2193624 | TPCN2 | 11 | 69048932 | 69136316 |
| VLDLR | 9 | 2621182 | 2660056 | DCT | 13 | 94436811 | 94479682 |
| SNX13 | 7 | 17790761 | 17940501 | ATP11A | 13 | 112690038 | 112887168 |
| GRM6 | 5 | 178977587 | 178996206 | SLC24A4 | 14 | 92322581 | 92501481 |
| ATF1 | 12 | 50763710 | 50821162 | KIAA0930 | 22 | 45192244 | 45240894 |
