## Extended Data Table 3 for "The multi-scale complexity of human genetic variation beyond continental groups"

| Type 2 Diabetes | | | |
|---|---|---|---|
| **Gene** | **Chromosome** | **Start (bp)** | **End(bp)** |
| HNF4A | 20 | 44355699 | 44432845 |
| RREB1 | 6 | 7107597 | 7251980 |
| GCKR | 2 | 27496839 | 27523684 |
| POC5 | 5 | 75674124 | 75717448 |
| ANKH | 5 | 14704800 | 14871778 |
| WSCD2 | 12 | 108129288 | 108250537 |
| KCNJ11 | 11 | 17365172 | 17389331 |
| PAM | 5 | 102753981 | 103029730 |
| TM6SF2 | 19 | 19264364 | 19273391 |
| LPL | 8 | 19901717 | 19967259 |
| PLCB3 | 11 | 64251530 | 64269150 |
| SLC30A8 | 8 | 116950273 | 117176714 |
| PNPLA3 | 22 | 43923792 | 43964488 |
| HNF1A | 12 | 120978543 | 121002512 |
| GIPR | 19 | 45668221 | 45683722 |
